# The third alpha helix of plant homeoproteins are generally cell-penetrating to plant cells

**DOI:** 10.1101/2025.04.01.646723

**Authors:** Henry J Squire, Jeffery W Wang, Markita P Landry

## Abstract

Homeoproteins are transcription factors involved in developmental regulation. Animal homeoproteins are generally cell-penetrating to animal cells, capable of crossing membranes through a conserved 3^rd^ alpha helix “penetration” domain. While plants also possess homeoproteins, the cell-penetrating ability of plant homeoproteins has not been investigated. We catalog the 3^rd^ alpha helixes of plant homeoproteins across 851 species and screen a subset for the ability to penetrate walled-plant cells. We identify plant 3^rd^ alpha helixes which are cell-penetrating to plants, the efficiency of which is amino acid sequence dependent, and use these peptides for cytosolic delivery of recombinant protein cargoes. We discover plant homeoproteins are generally cell-penetrating to plants, mirroring the behavior of animal homeoproteins, with applications in plant biotechnology and implications for fundamental plant biology.

## Main Text

Homeoproteins are DNA binding factors conserved across eukaryotes and are typically involved in developmental regulation (*1*). First discovered in *Drosophila* (*2*), homeoproteins are encoded by homeobox genes and are classified as homeoproteins on the basis of a highly conserved 60 amino acid sequence consisting of three consecutive alpha helixes. Homeobox genes have been identified across many taxa of life including vertebrates (*3*) which typically possess hundreds of unique homeoproteins (*1*). In 1991, Joliot *et al*. reported the *Drosophila* homeoprotein Antennapedia was capable of “self” internalizing into neuronal cells, independently translocating across biological membranes into nuclei (*4, 5*). Follow up work identified the 3^rd^ alpha helix of the homeodomain of Antennapedia as responsible for this unexpected behavior (*6*). This peptide sequence became known as Penetratin and is the second ever discovered cell-penetrating peptide (CPP). Since this discovery, animal homeoproteins have been found to be generally cell-penetrating to animal cells (*7, 8*), the role of animal homeoproteins’ ability to translocate across cells in developmental regulation has been explored (*9*), and the cell-penetrating capability of Penetratin (as well as other CPPs) has been exploited to deliver cargoes such as DNA, RNA, protein, and drugs to animal cells for clinical applications (*10, 11*).

To date, the field of CPPs has focused primarily on peptides derived from non-plant sources with a focus on mammalian cell and clinical applications. Plants, like animals, also possess homeoproteins, with WUSCHEL (WUS), SHOOT MERISTEMLESS (STM), and KNOTTED1 (KN1) a few of the best studied examples. Previous work demonstrated the 3^rd^ alpha helix of the plant homeoprotein KNT1 is cell-penetrating to animal cells (*12*). Separate work demonstrated KNT1 and STM traffic between plant cells through plasmodesmata acting non-cell autonomously (*13, 14*), a phenomena dependent specifically on the homeodomain. This prior work hinted plant homeoproteins might be cell-penetrating to plants, similar to the manner in which animal homeoproteins are cell-penetrating to animals. However, thus far the cell-penetrating ability of plant homeoproteins to plants has not been investigated. We specifically hypothesized the 3^rd^ alpha helixes from the homeodomains of plant homeoproteins are generally cell-penetrating to plants. To test this hypothesis, we cataloged the 3^rd^ alpha helixes of plant homeoproteins by mining the UniProtKB database. We next screened a subset of plant 3^rd^ alpha helix peptides for the ability to penetrate plant cells (cellular internalization to the cytosol) using a previously reported fluorescence complementation method termed delivered complementation in-planta (DCIP) (*15*). Through this approach, we demonstrate the 3^rd^ alpha helix of plant homeoproteins are generally cell-penetrating to plants with applications in applied plant biotechnology and implications for fundamental plant biology.

### WUSCHEL homeoprotein self-internalizes into plant cells

In our previous work, we developed a split GFP complementation method termed delivered complementation in planta (DCIP) for quantification of direct exogenous delivery of proteins or peptides to plants (*15*). DCIP relies on expression of a nuclear localized fusion protein of sfGFP1-10 and mCherry in the leaves of *Nicotiana benthamiana*, through *A. tumefasciens* infection, followed by syringe infiltration of an exogenous protein or peptide tagged with GFP11 (Fig. 1A). Both sfGFP1-10 and GFP11 lack fluorescence independently (*16*) but become fluorescent upon spontaneous complementation of the two components. Complementation can only occur if GFP11 peptide successfully internalizes to the plant cell with the assistance of a delivery tool such as a CPP. Infiltrated leaves are imaged with laser scanning confocal microscopy and the cellular internalization efficiency of the exogenously applied protein or peptide is calculated by the ratio of the number of nuclei with GFP signal to the number of nuclei with mCherry signal (% GFP positive nuclei).

**Fig. 1.**
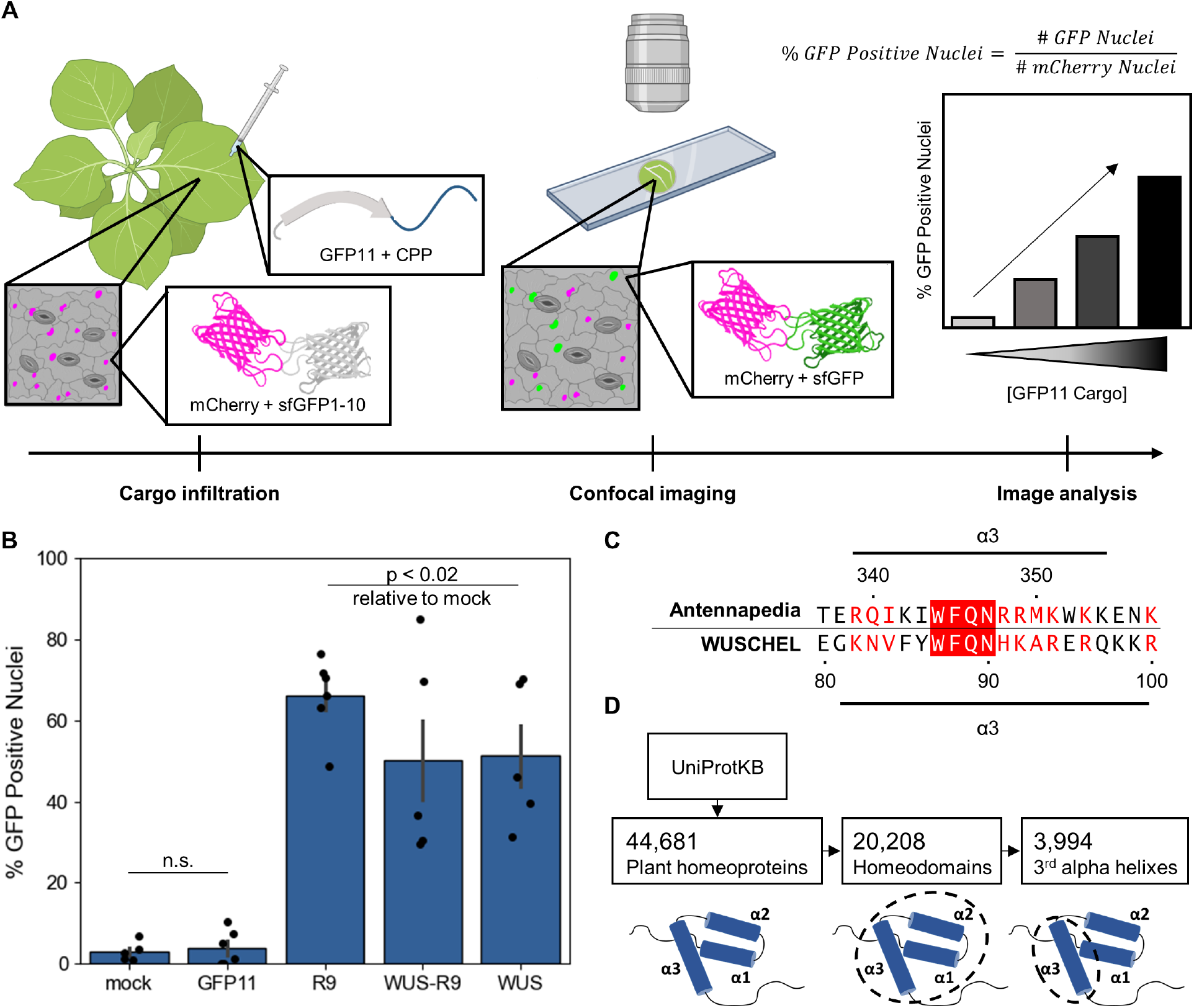
DCIP work flow, validation, and mining of plant homeoproteins. **A)** Leaves of four-week old *Nicotiana benthamiana* leaves are transfected with *A. tumefasciens* to express a NLS-mCherry-GFP1-10 fusion protein. Three days later, leaves are syringe infiltrated with CPP fused to GFP11. Five hours later, leaf punches are imaged with confocal microscopy. Using automated image analysis, the number of nuclei expressing GFP and/or mCherry are counted. The ratio of the number of GFP nuclei to the number of mCherry nuclei represents the cellular internalization efficiency of the exogenously applied peptide or protein. **B)** Bar graph of average % GFP positive nuclei in *N. benthamiana* leaves infiltrated with: water (mock), 100 μM GFP11 (GFP11), 100 μM R9 CPP fused to GFP11 (R9), 140 μM of recombinant WUS with a N-terminal GFP11 tag and C-terminal R9 CPP (WUS-R9), and 140 μM of recombinant WUS with a N-terminal GFP11 tag without a C-terminal R9 CPP (WUS). Each point represents a biological replicate from a single leaf derived from the average of 4 fields of view. Bars represent the average of at least 5 biological replicates. Kruskal-Wallis test followed by Dunn’s multiple comparison post-hoc was used to compare % GFP positive nuclei of each treatment to mock treatment. Data taken from (*15*). **C)** Alignment of the 3^rd^ alpha helix from the homeodomain of Antennapedia (Penetratin) and the 3^rd^ alpha helix from the homeodomain of WUSCHEL. Aligned residues with chemical similarity are marked in red while aligned residues with exact identity matches are marked in white with a red outline. **D)** Illustration of our database mining approach to identify 3^rd^ alpha helix peptides from the homeodomain of plant homeoproteins. Boxed numbers represent the number of unique sequences identified at each step.

We leveraged DCIP to test exogenous delivery of proteins to *N. benthamiana* using CPPs. First, we demonstrated nona-arginine (R9), a model CPP from animal literature, tagged with GFP11 efficiently internalized to leaf cells (Fig. 1B, data from (*15*)). Next we sought to deliver a full recombinant protein with R9; we translationally fused GFP11 and a R9 CPP to the N and C termini respectively of recombinant WUSCHEL (WUS) protein. DCIP confirmed successful cellular internalization of the recombinant WUS construct in *N. benthamiana* leaves (Fig. 1B, data from (*15*)). In a control experiment, we produced a recombinant WUS protein without the C terminal R9 CPP and we unexpectedly found WUS still internalized at the same efficiency as WUS-R9 (Fig. 1B, data from (*15*)). GFP11 alone, without a R9 CPP, did not internalize, suggesting a domain of WUS was enabling cellular internalization.

We hypothesized the homeodomain of WUS, spSecifically the 3^rd^ alpha helix, was responsible for the unexpected ability of WUS to self-internalize without a CPP. We aligned the 3^rd^ alpha helix of the homeodomain of Antennapedia (also called Penetratin, the second ever discovered CPP) (*6*) to the 3^rd^ alpha helix of the homeodomain of WUS and found high sequence similarity (Fig. 1C), supporting our hypothesis. We further hypothesized the 3^rd^ alpha helix of plant homeoproteins might be generally cell-penetrating to plants.

To probe this hypothesis, we first explored the landscape of plant homeoproteins from which to derive 3^rd^ alpha helix peptides. We consulted the UniProtKB (*17*) database (data S1) identifying a total of 40,351 plant homeoproteins (Fig. 1D and data S2) from 851 diverse plant species (fig. S1). We extracted the homeodomain (typically 60 amino acids in length) from each plant homeoprotein, identifying 20,208 unique homeodomains and aligned the homeodomains identifying 3,994 unique 3^rd^ alpha helix peptides (Fig. 1D). Certain positions along the 3^rd^ alpha helix were highly conserved, most notably a tryptophan at position 6 and a phenylalanine at position 7 found in 95% and 88% of 3^rd^ alpha helix peptides respectively (fig. S2) in agreement with previous work, validating our classification of these peptides as 3^rd^ alpha helixes (*18*). To investigate if the 3^rd^ alpha helix of plant homeoproteins are generally cell-penetrating, we selected 30 for experimentation. These 3^rd^ alpha helix peptides were drawn from a diverse set of plant species representing all 14 classes of plant homeoproteins (*18*). A list of 3^rd^ alpha helix peptides selected for experimentation can be found in data S3 along with a sequence alignment in fig. S3.

### Internalization of 3^rd^ alpha helix peptides into plant cells

Selected 3^rd^ alpha helix peptides were subsequently screened for the ability to internalize to plants at a concentration of 50 μM using DICP. Representative maximum intensity projections from DCIP experiments in *N. benthamiana* leaves are shown in Fig. 2A (overlay with chloroplast autofluorescence in fig. S4). In general, 3^rd^ alpha helix peptides derived from plant homeoproteins are cell-penetrating to *N. benthamiana* leaves; 21 of 30 peptides internalized at a significantly greater efficiency than mock treatment (Fig. 2B and fig S5). As a control, a non-homeoprotein, CLSY2 (*19*), was aligned against the 3^rd^ alpha helix peptides; this non-homeoprotein derived peptide was screened with DCIP, and found to be not cell-penetrating (labeled “random” in Fig. 2B). To control for potential interference of the 3^rd^ alpha helix peptide with complementation between GFP11 and GFP1-10, an *in vitro* complementation assay was conducted. With the exception of the peptides GgBEL and StPHD, all other peptides complemented efficiently *in vitro* (fig. S6) suggesting differences in the percentage of GFP positive nuclei between peptides are due to discrepancies in cellular internalization efficiency not fluorescence complementation efficiency.

**Fig. 2.**
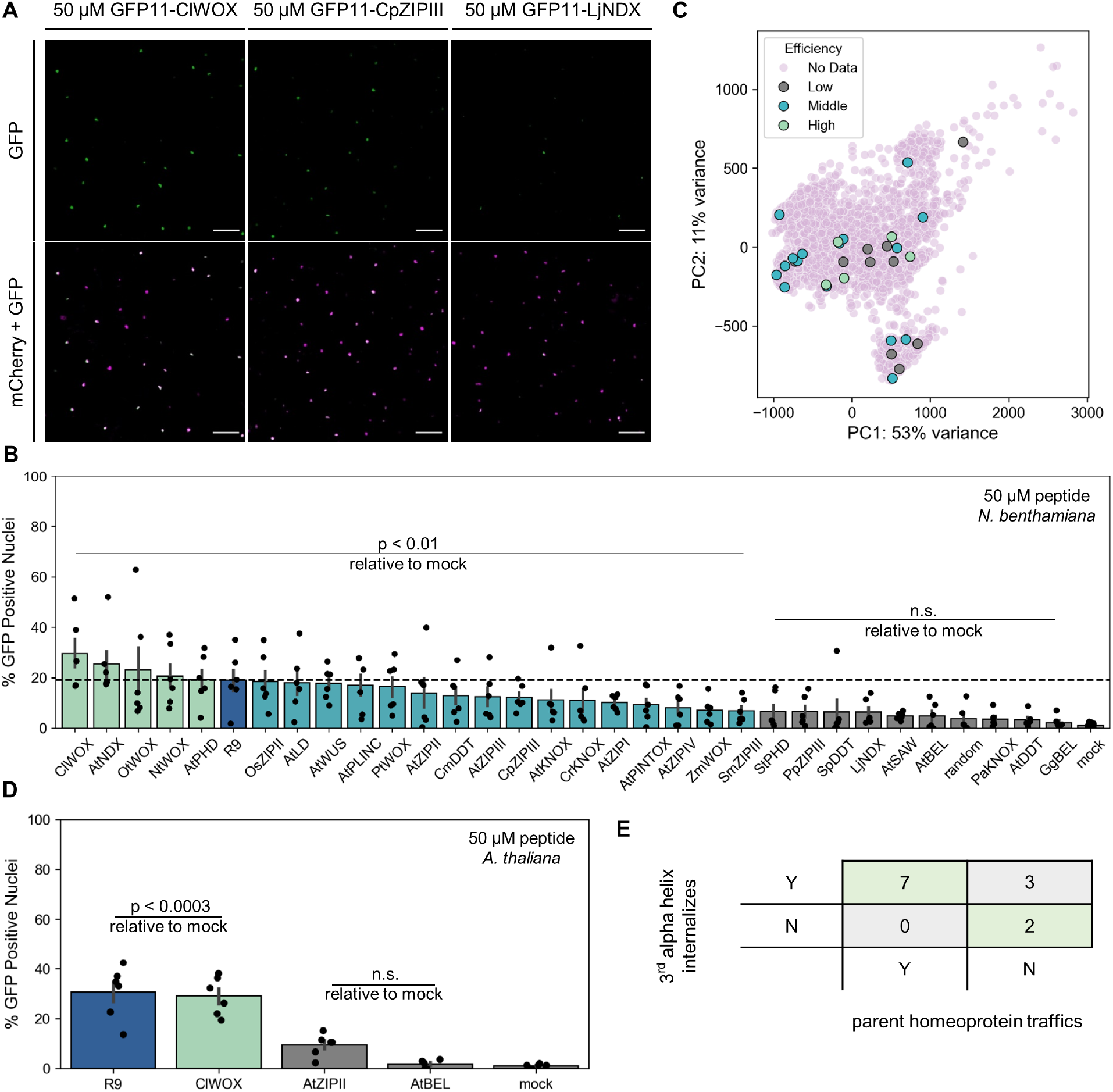
The 3^rd^ alpha helix of plant homeoproteins is cell-penetrating. **A)** Representative maximum intensity projections of *N. benthamiana* leaves expressing the DCIP reporter system infiltrated with 50 μM of different 3^rd^ alpha helix peptides fused to GPF11. The first column, displays leaves infiltrated with ClWOX, a peptide with high internalization efficiency. The second column displays leaves infiltrated with CpZIPIII, a peptide with medium internalization efficiency. The third column displays leaves infiltrated with LjNDX, a peptide with little to no internalization efficiency. The top row displays GFP signal pseudo-colored green. The bottom row displays a two-color overlay of mCherry signal pseudo-colored magenta with GFP. Colocalization of GFP and mCherry results in white in the two-color overlay in the bottom row. Scale bars represent 100 μm. **B)** Bar graph of the average % GFP positive nuclei for *N. benthamiana* leaves infiltrated with 50 μM of different 3^rd^ alpha helix peptides fused to GPF11. A model CPP, R9, is included for comparison, marked in blue with the average % GFP positive nuclei for R9 marked with a dashed line. Sequences with an average % GFP positive nuclei greater than R9 are colored green, sequences with an average % GFP positive nuclei less than R9 but statistically significantly greater than water mock treatment are colored teal, and sequences with an average % GFP positive nuclei statistically indistinguishable from water mock treatment are colored gray. Each point represents a biological replicate derived from four fields of view taken from a single leaf with a total of six different leaves drawn from distinct plants. Error bars represent the standard error of the mean. Kruskal-Wallis test followed by Dunn’s multiple comparison post-hoc was used to compare % GFP positive nuclei of each peptide to mock treatment. **C)** Scatter plot of the first two principal components from a principal component analysis on a BLOSUM62 substitution of all 3,994 plant 3^rd^ alpha helix peptides identified here. The 3^rd^ alpha helix peptides which were experimentally screened for cellular internalization efficiency in *N. benthamiana* leaves are overlaid and color coded by efficiency. **D)** Bar graph of the average % GFP positive nuclei for the leaves of *A. thaliana* seedlings treated with 50 μM of different 3^rd^ alpha helix peptides fused to GPF11. A model CPP, R9, is included for comparison, marked in blue. Each point represents a biological replicate derived from the average of three fields of view taken from a single seedling with a total of six different seedlings. Error bars represent the standard error of the mean. Kruskal-Wallis test followed by Dunn’s multiple comparison post-hoc was used to compare % GFP positive nuclei of each peptide to mock treatment. **E)** Correlation between a parent plant homeoprotein’s ability to traffic and the ability of the corresponding 3^rd^ alpha helix for cellular internalization. “Y” indicates ability to traffic or internalize while “N” indicates lack of ability to traffic or internalize.

Cellular internalization efficiency at 50 μM is highly variant, as high as 30% for the 3^rd^ alpha helix peptide ClWOX to as low as 2% for the 3^rd^ alpha helix peptide GgBEL. Cellular internalization efficiency does not specifically depend on the species or homeodomain class from which the 3^rd^ alpha helix peptide was derived. Furthermore, a principal component analysis of a BLOSUM62 substitution of all 3^rd^ alpha helix peptides shows peptides do not cluster by cellular internalization efficiency (Fig. 2C). Compared to the model CPP R9 (marked in dark blue in Fig. 2B), five 3^rd^ alpha helix peptides internalize on average as efficiently or slightly more efficiently (marked in green in Fig. 2B). R9 is currently amongst one of the most efficient CPPs in plants (*15*); 3^rd^ alpha helix peptides with even greater cellular internalization efficiencies likely exist in the 3,994 unique 3^rd^ alpha helix peptides cataloged here.

Separately, we screened three 3^rd^ alpha helix peptides, ClWOX, AtZIPII, and AtBEL, in transgenic *Arabidopsis thaliana* expressing DCIP to evaluate the species dependency of our findings. Similar to *N. benthamiana* leaves, ClWOX and R9 are both cell-penetrating to the leaves of *A. thaliana* seedlings (Fig. 2D and fig. S7 & S8). AtZIPII and AtBEL, in contrast, are not cell-penetrating with cellular internalization efficiencies indistinguishable from mock treatment (Fig. 2D). As with *N. benthamiana*, ClWOX, has the highest cellular internalization efficiency of the investigated 3^rd^ alpha helix peptides. These results suggest the 3^rd^ alpha helix of plant homeoproteins are cell-penetrating to plants generally, not just *N. benthamiana* specifically.

Interestingly, the internalization ability of the 3^rd^ alpha helix peptides correlates with the endogenous cell-to-cell trafficking ability of the parent homeoprotein from which the 3^rd^ alpha helix peptide was derived. Here, trafficking refers to a phenomenon in which homeoproteins are found in cells which lack the corresponding coding mRNA. For example, the 3^rd^ alpha helix peptides AtKNOX and AtWUS have cellular internalization efficiencies in *N. benthamiana* of 11% and 18% respectively (Fig. 2B) and are both derived from homeoproteins, KN1 and WUS respectively, which are known to traffic (*14, 20*). In contrast, the internalization efficiency of the peptide AtBEL is indistinguishable from mock treatment, effectively not cell-penetrating (Fig. 2C), and is derived from a homeoprotein, BELLRINGER, which is known to not traffic (*20*).

Data on endogenous plant homeoprotein trafficking is limited compared to animal homeoprotein trafficking; from our literature search, we identified 14 plant homeoproteins which were explicitly studied for trafficking ability. For reference, data on trafficking of at least 170 animal homeoproteins is available (*7*). We identified seven plant homeoproteins, reported in literature to endogenously traffic (table S1), with their corresponding 3^rd^ alpha helix internalizing and two plant homeoproteins, reported in literature to not endogenously traffic (table S1), with their corresponding 3^rd^ alpha helix not internalizing (Fig. 2E). We identified three plant homeoproteins, reported in literature to not endogenously traffic (table S1) with their corresponding 3^rd^ alpha helix still internalizing. Our results suggest there is a correlation between endogenous homeoprotein trafficking and exogenous 3^rd^ alpha helix internalization ability. The biological implications of this correlation is not clear and additional data on plant homeoprotein trafficking is necessary, though we hypothesize apoplastic trafficking of homeoproteins mediated by the cell-penetrating capability of the 3^rd^ alpha helix could explain this correlation. As such, our discovery that the 3^rd^ alpha helix of plant homeoproteins are cell-penetrating may have implications beyond plant biotechnology in fundamental plant biology.

### Mechanisms of 3^rd^ alpha helix internalization to plant cells

We next explored why certain 3^rd^ alpha helix peptides efficiently internalize in *N. benthamiana* leaves. We first note minor changes in amino acid sequence drastically impact cellular internalization efficiency. For example, the 3^rd^ alpha helix peptides AtNDX and LjNDX have similar amino acid compositions, different by only two amino acids at positions 13 and 17, yet AtNDX is cell-penetrating with a cellular internalization efficiency of 26% in *N. benthamiana* while LjNDX is not appreciably cell-penetrating, indistinguishable from mock treatment (Fig 3A). Exploring the positions with differences more closely, at position 13, AtNDX and LjNDX bear arginine and lysine residues respectively, amino acids which are chemically similar (both polar positive). Meanwhile, at position 17, AtNDX and LjNDX bear an alanine and threonine residue respectively, chemically different residues (hydrophobic versus polar). The only true chemical difference between AtNDX and LjNDX is at position 17; hence, a single residue substitution can significantly impact the ability of a 3^rd^ alpha helix peptide to internalize. Similar observations can be made with NtWOX and ZmWOX as well as ClWOX and PtWOX (Fig. 3A), pointing to the complexity of the relationship between amino acid sequence and 3^rd^ alpha helix peptide internalization.

**Fig 3.**
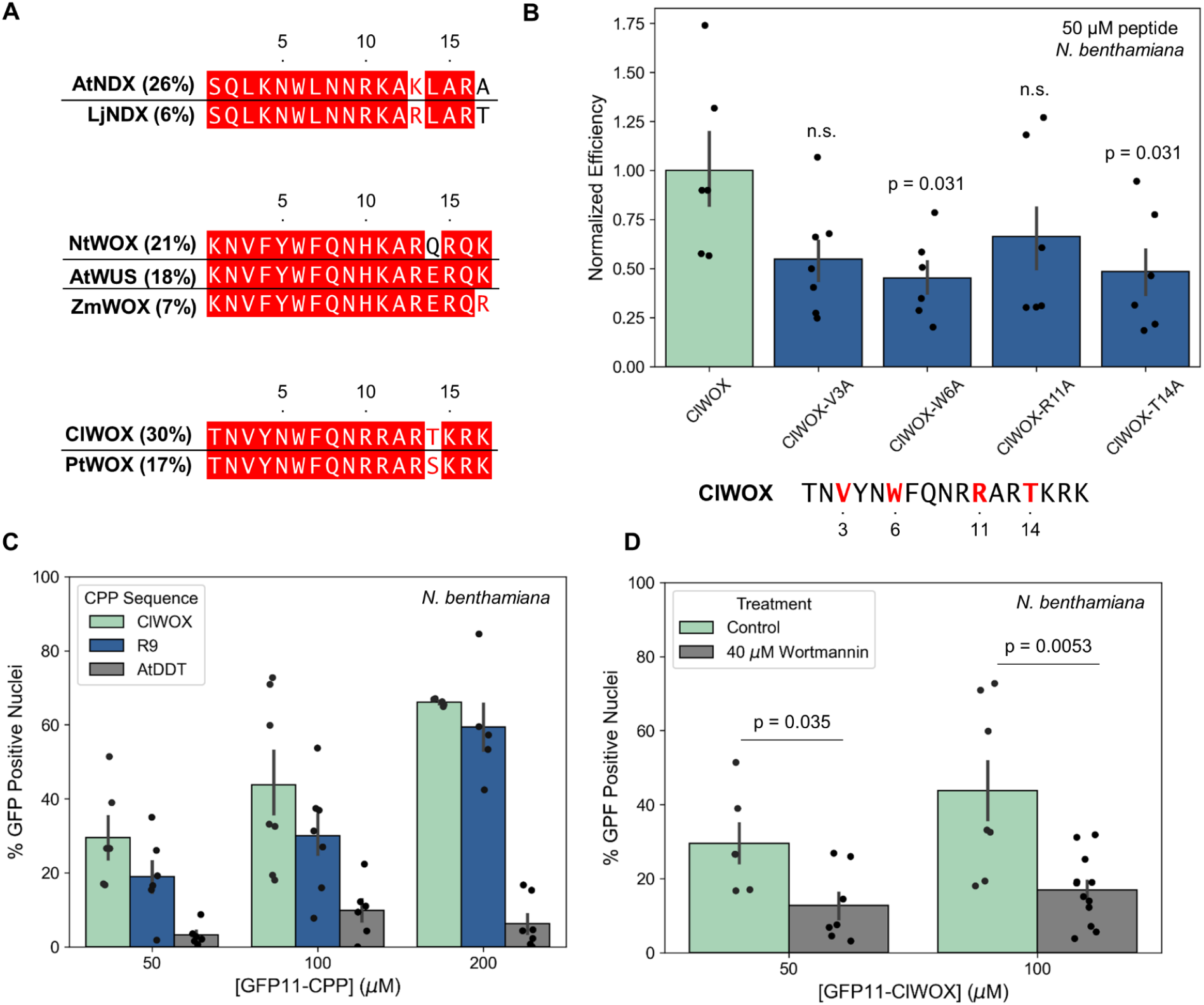
Conditions and mechanisms influencing 3^rd^ alpha helix internalization. **A)** Sequence alignment between select 3^rd^ alpha helix peptides with similar amino acid composition but different cellular internalization efficiency. Aligned residues with exact identity matches are marked in white with a red outline, aligned residues with chemical similarity are marked in red, and aligned residues which are chemically divergent are marked in black. The average cellular internalization efficiency of each 3^rd^ alpha helix peptide is marked in parenthesis. **B)** Bar graph displaying average cellular internalization efficiency for different alanine mutated ClWOX peptides normalized to the internalization efficiency of non-mutated ClWOX peptide in the leaves of *N. benthamiana*. Peptides were tested at a concentration of 50 μM. Each point represents the average of four fields of view taken from a single leaf with a total of six different leaves drawn from distinct plants. Error bars represent the standard error of the mean. Kruskal-Wallis test followed by Dunn’s multiple comparison post-hoc was used for pair wise comparison of the normalized efficiency of each mutated peptide to ClWOX. The p-value is reported above each bar, n.s. indicates p > 0.05. Non-mutated ClWOX is shown below the bar graph and positions which were mutated to alanine are marked in red. Only one position was mutated for each peptide tested. **C)** Bar graph of the average % GFP positive nuclei for different 3^rd^ alpha helix peptides at 50 μM, 100 μM, and 200 μM concentrations in the leaves of *N. benthamiana*. Each point represents four fields of view taken from a single leaf with a total of three to seven different leaves drawn from distinct plants tested for each peptide. Error bars represent the standard error of the mean. Kruskal-Wallis test followed by Dunn’s multiple comparison post-hoc was used for pair wise comparison of each peptide and each concentration in fig. S9. **D)** Bar graph of the average % GFP positive nuclei at different concentrations of ClWOX with and without pretreatment with 40 μM of Wortmannin endocytosis inhibitor in the leaves of *N. benthamiana*. Each point represents the average of four fields of view taken from a single leaf with a total of six to eleven different leaves (drawn from distinct plants) tested for each condition. Error bars represent the standard error of the mean. Kruskal-Wallis test was used for pair wise comparison of Wortmannin pre-treated leaves to control leaves at each respective ClWOX concentration.

To further investigate this relationship, we conducted a partial alanine mutagenesis screen on the most efficient 3^rd^ alpha helix peptide, ClWOX, focusing on previously reported amino acid residues which are critical for internalization of Penetratin to animal cells (*21*). All alanine mutations reduced cellular internalization efficiency in *N. benthamiana* (Fig. 3B). Mutating tryptophan at position 6 most significantly reduced internalization with delivery efficiency reduced by half. Previous work on Penetratin similarly found the tryptophan in the hydrophobic core of the peptide critical for internalization (*21*). The only other peptide investigated here which lacks a tryptophan at position 6 is AtLD which instead bears a phenylalanine (fig. S3). This suggests cyclic hydrophobic amino acids at position 6 are important though not sufficient for efficient internalization of a 3^rd^ alpha helix peptide in *N. benthamiana* leaves. Identifying clear rules to explain internalization is not trivial due to the complex and surprisingly sensitive relationship between amino acid sequence and 3^rd^ alpha helix peptide internalization efficiency.

Next, we investigated the concentration dependence of cellular internalization efficiency selecting ClWOX and AtDDT, representing amongst the highest and lowest efficiency 3^rd^ alpha helix peptides respectively. We increased the concentrations which we infiltrated to *N. benthamiana* to 100 μM and 200 μM. For ClWOX and R9, cellular internalization efficiency is concentration dependent with a maximum efficiency of 66% for ClWOX at 200 μM (Fig. 3C). For AtDDT, internalization efficiency increased slightly with concentration, yet still not statistically distinguishable from mock treatment. Net, cellular internalization efficiency is dependent on amino acid sequence as well as concentration.

We next probed the mechanism of uptake of the high efficiency ClWOX peptide at 50 μM and 100 μM concentrations. CPPs internalize through energy independent and/or energy dependent pathways in animal cells depending on the cell type, cargo, CPP concentration, and CPP amino acid sequence (*22*). We pretreated *N. benthamiana* leaves with 40 μM of Wortmannin, an endocytosis inhibitor (*23*), to generically block/reduce energy dependent uptake and infiltrated leaves with ClWOX peptide forty-five minutes later. Average cellular internalization efficiency was reduced by pre-treatment with Wortmannin but not totally abolished across both concentrations of ClWOX (Fig. 3D). This data suggests both energy independent and energy dependent pathways play a role in internalization. In animal cells, CPPs typically internalize through a mix of energy dependent and energy independent pathways simultaneously (*22*). Our results are broadly in line with this body of work.

### Functional cargo delivery via a plant 3^rd^ alpha helix peptide

Finally, to test the utility of 3^rd^ alpha helix peptides in plant biotechnology, we attempted to deliver a functional protein cargo to walled plant cells. We sought to deliver Cre recombinase, an enzyme which catalyzes DNA recombination with application in plant biotechnology to excise for example selectable marker genes in production of transgenic plants (*24*). To this end, we translationally fused ClWOX to the C-terminus of Cre recombinase and designed a reporter consisting of two transcriptional units driving either mCherry or GFP expression. GFP expression was blocked by a premature terminator flanked by loxP sites such that successful delivery of functional Cre recombinase would excise the premature terminator and turn on expression of GFP (Fig. 4A). *N. benthamiana* leaves were transfected with *A. tumefasciens* to express the reporter system and syringe infiltrated with 10 μM of recombinant Cre-ClWOX fusion two days after transfection. Confocal microscopy 24 hours after infiltration confirmed successful delivery of Cre recombinase, indicated by GFP expression in *N. benthamiana* leaves (Fig. 4B and fig. S10). On average, 19% of cells expressed GFP post-infiltration with 10 μM Cre-ClWOX (Fig. 4C). Given the dose dependence observed in Fig. 3C, we expect Cre recombinase delivery efficiency would increase with concentration. Cre recombinase without a 3^rd^ alpha helix tag did not successfully turn on GFP expression. The 3^rd^ alpha helix of plant homeoproteins can be used to deliver functional recombinant proteins to walled plant cells for plant biotechnology applications.

**Fig 4.**
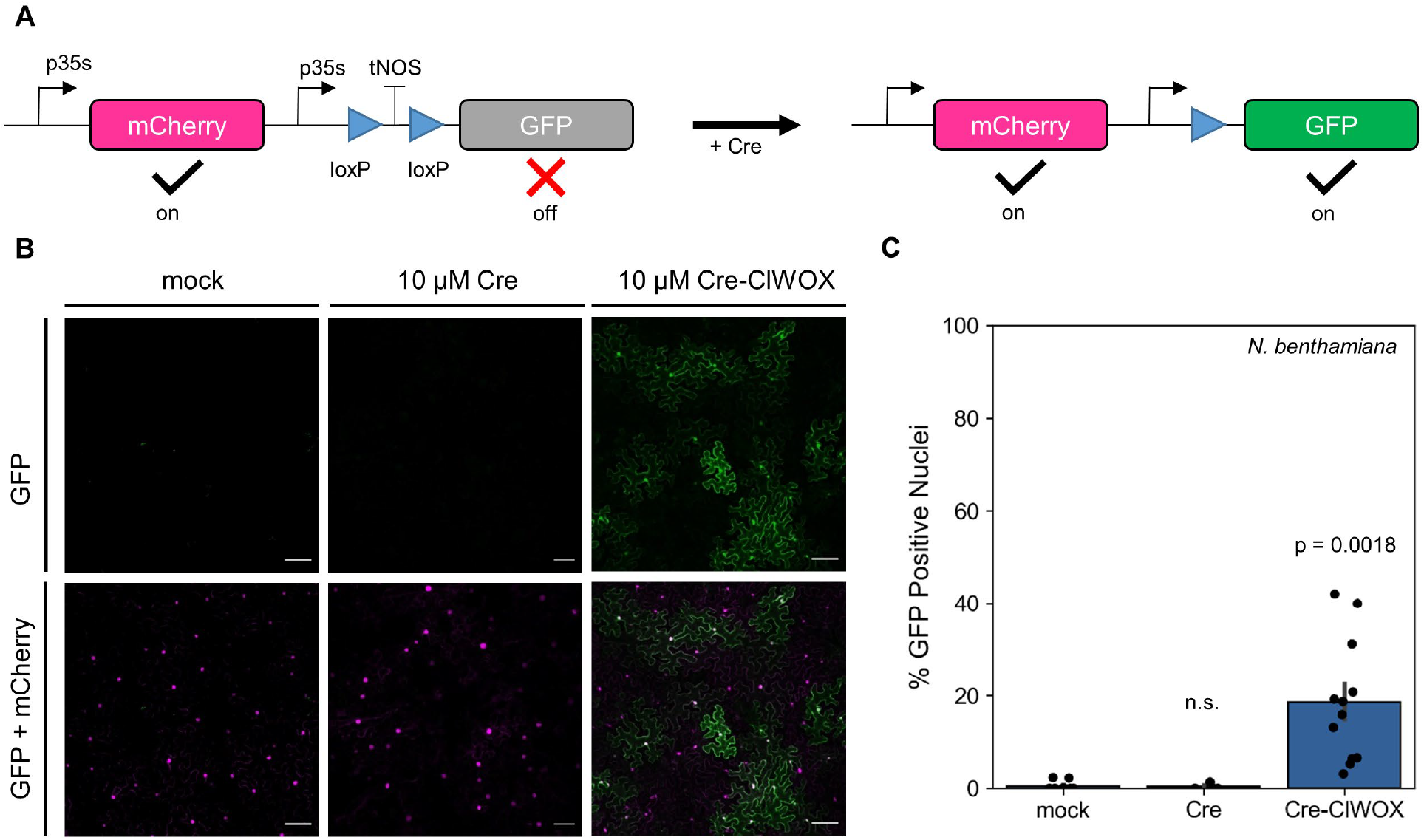
Delivery of Cre recombinase via 3^rd^ alpha helix peptide. **A)** Schematic of Cre recombinase reporter system consisting of two transcriptional units, one which drives expression of a nuclear localized mCherry protein and one which drives expression of nuclear localized GFP. While mCherry is constantly expressed (on), expression of GFP is blocked by a premature terminator flanked by loxP sites (off). Exogenous delivery of functional Cre recombinase results in excision of the premature terminator resulting in expression of GFP as well as mCherry. **B)** Representative maximum intensity projections of *N. benthamiana* leaves expressing the Cre recombinase reporter system 24 hours after mock, 10 μM Cre (no CPP), or 10 μM Cre-ClWOX treatment. The first row displays GFP signal pseudo colored green. The second row displays two color overlay of mCherry signal (pseudo colored magenta) and GFP signal. Colocalization of GFP and mCherry appears as white in the two color overlay. All scale bars represent 100 μm. **C)** Quantification of maximum intensity projections displaying the average % GFP positive nuclei for *N. benthamiana* leaves expressing the Cre recombinase reporter for different treatments. Each point represents a single leaf drawn from distinct plants with at least 4 leaves tested per treatment. Error bars represent the standard error of the mean. Kruskal-Wallis test was used for pair wise comparison of different Cre treatments to mock treatment.

## Discussion

Direct delivery of protein to walled plant cells remains a challenge. While a variety of methods are available to directly deliver protein cargoes to animal cells, many of these methods are ineffective or unexplored in plants. CPPs in particular are well-studied in animals with fewer studies in plants. While over 1,800 unique CPP sequences are reported in animal literature (*25*), only a small subset have been studied in plants (*26*). Here we demonstrate the 3^rd^ alpha helix of the homeodomain of plant homeoproteins serve as a source of novel CPP sequences for efficiently delivering cargoes to plant cells. We selected 3^rd^ alpha helixes for experimentation on the basis of diversity in amino acid sequence, species of origin, and classification, without prior knowledge of cellular internalization efficiency. Without explicitly searching for efficient CPPs, five 3^rd^ alpha helix peptides with average cellular internalization efficiencies equivalent to R9 were identified. The relationship between peptide amino acid sequence and delivery efficiency is complicated, a single mutation in an efficient peptide can render the peptide non-functional. Uncovering the relationship between amino acid sequence and cellular internalization efficiency could enable mining of plant homeoproteins for CPPs with even higher cellular internalization efficiency.

Numerous studies have attempted to elucidate the mechanisms by which CPPs internalize to animal cells. However, inquiry into the mechanisms of CPP internalization to plants cells is limited owing to historic the lack of tools to confirm protein delivery (*15, 27*). Given the 3^rd^ alpha helix peptides investigated here appear to internalize via a mix of energy dependent and energy independent processes, endosomal escape might represent a potential bottleneck in achieving efficient cytosolic delivery of proteins in plants. In animal cells, endosomal escape post energy dependent uptake (endocytosis) is known to be inefficient (*28*), yet in plants, the efficiency of endosomal escape remains uncharacterized.

In animal cells, extracellular trafficking of homeoproteins is critical in developmental regulation. In contrast, protein trafficking occurs through plasmodesmata in plants (*14*). Homeoprotein trafficking in animal cells depends on a cellular secretion domain and cellular penetration domain (*7, 29*); typically, the secretion domain is found in between the 2^nd^ and 3^rd^ alpha helix of the homeodomain while the penetration domain is found in the 3^rd^ alpha helix of the homeodomain (*30*). To our knowledge, similar secretion and penetration domains have not been identified in plant homeoproteins; our work here identifies a penetration domain. Previous work demonstrated the homeodomain is involved in the ability of plant homeoproteins to traffic endogenously in plants (*31*). In the plant homeoprotein KN1 for example, the 3^rd^ alpha helix of the homeodomain is necessary to enable cell-to-cell trafficking with the whole homeodomain necessary and sufficient for trafficking (*20*). Similarly, the homeodomain of WUS promotes trafficking (*32*). Notably, not all plant homeoproteins traffic (*14*) and we observe some correlation between a plant homeoprotein’s ability to traffic and the ability of its 3^rd^ alpha helix to internalize into plant cells. We suspect secretion and extracellular trafficking is an unlikely mode of homeoprotein transport in plants due to plasmodesmata and the cell wall, although we do not rule out the possibility of secretion and extracellular trafficking via apoplastic pathways. Taken together, our results present plant homeoproteins, specifically the 3^rd^ alpha helix of the homeodomain, as a new tool for protein delivery with applications in plant biotechnology and implications in plant biology.

## Acknowledgments

Confocal microscopy experiments were conducted at the CRL Molecular Imaging Center, RRID:SCR_017852, supported by the Helen Willis Neuroscience Institute. We thank Holly Aaron and Feather Ives for discussions about imaging. We thank Eyal Fridman for discussions about protein purification. We thank Ruby Nelson and Jamie Phan for maintaining plants for this work. We thank Karthik Shekhar and Ben Williams for insightful discussion and feedback on this work.

## Funding

Burrough Wellcome Fund Career Award at the Scientific Interface (MPL)

Dreyfus foundation award (MPL)

Philomathia foundation (MPL)

NSF CAREER award 2046159 (MPL)

McKnight Foundation award (MPL)

Simons Foundation Award (MPL)

Moore Foundation Award (MPL)

Heising-Simons Fellowship (MPL)

Brain Foundation Award (MPL)

polymaths award from Schmidt Sciences, LLC (MPL)

National Defense Science and Engineering Graduate Fellowship U.S. Department of Defense (HJS)

Philomathia Foundation Graduate Fellowship (HJS)

## Author contributions

Conceptualization: HJS, JWW, MPL

Methodology: HJS, JWW

Investigation: HJS

Visualization: HJS

Funding acquisition: MPL

Project administration: MPL

Supervision: MPL

Writing – original draft: HJS

Writing – review & editing: HJS, JWW, MPL

## Data and materials availability

All data necessary to evaluate this work are available in the main text, supplementary material, or supplementary data. Underlying source images and code for analysis for all figures are available in Dryad. Further correspondence or material/data requests should be addressed to MPL.

## Supplementary Materials

Material and Methods

Figs. S1 to Figure S13

Table S1

References (*33–37*)

## Other Supplementary Data

Data S1 to S3

## References and Notes

1. T. R. Bürglin, M. Affolter, Homeodomain proteins: an update. Chromosoma 125, 497–521 (2016).

2. W. McGinnis, R. L. Garber, J. Wirz, A. Kuroiwa, W. J. Gehring, A homologous protein-coding sequence in drosophila homeotic genes and its conservation in other metazoans. Cell 37, 403–408 (1984).

3. W. McGinnis, M. S. Levine, E. Hafen, A. Kuroiwa, W. J. Gehring, A conserved DNA sequence in homoeotic genes of the Drosophila Antennapedia and bithorax complexes. Nature 308, 428–433 (1984).

4. A. H. Joliot, A. Triller, M. Volovitch, C. Pernelle, A. Prochiantz, alpha-2,8-Polysialic acid is the neuronal surface receptor of antennapedia homeobox peptide. New Biol 3, 1121–1134 (1991).

5. A. Joliot, C. Pernelle, H. Deagostini-Bazin, A. Prochiantz, Antennapedia homeobox peptide regulates neural morphogenesis. Proc Natl Acad Sci U S A 88, 1864–1868 (1991).

6. D. Derossi, A. H. Joliot, G. Chassaing, A. Prochiantz, The third helix of the Antennapedia homeodomain translocates through biological membranes. Journal of Biological Chemistry 269, 10444–10450 (1994).

7. E. J. Lee, N. Kim, J. W. Park, K. H. Kang, W. Kim, N. S. Sim, C.-S. Jeong, S. Blackshaw, M. Vidal, S.-O. Huh, D. Kim, J. H. Lee, J. W. Kim, Global Analysis of Intercellular Homeodomain Protein Transfer. Cell Reports 28, 712-722.e3 (2019).

8. S. Balayssac, F. Burlina, O. Convert, G. Bolbach, G. Chassaing, O. Lequin, Comparison of Penetratin and Other Homeodomain-Derived Cell-Penetrating Peptides: Interaction in a Membrane-Mimicking Environment and Cellular Uptake Efficiency. Biochemistry 45, 1408–1420 (2006).

9. J. Apulei, N. Kim, D. Testa, J. Ribot, D. Morizet, C. Bernard, L. Jourdren, C. Blugeon, A. A. Di Nardo, A. Prochiantz, Non-cell Autonomous OTX2 Homeoprotein Regulates Visual Cortex Plasticity Through Gadd45b/g. Cerebral Cortex 29, 2384–2395 (2019).

10. M. P. Schutze-Redelmeier, H. Gournier, F. Garcia-Pons, M. Moussa, A. H. Joliot, M. Volovitch, A. Prochiantz, F. A. Lemonnier, Introduction of exogenous antigens into the MHC class I processing and presentation pathway by Drosophila antennapedia homeodomain primes cytotoxic T cells in vivo. J Immunol 157, 650–655 (1996).

11. D. Derossi, G. Chassaing, A. Prochiantz, Trojan peptides: the penetratin system for intracellular delivery. Trends in Cell Biology 8, 84–87 (1998).

12. M. Tassetto, A. Maizel, J. Osorio, A. Joliot, Plant and animal homeodomains use convergent mechanisms for intercellular transfer. EMBO Reports 6, 885–890 (2005).

13. D. Jackson, B. Veit, S. Hake, Expression of maize KNOTTED1 related homeobox genes in the shoot apical meristem predicts patterns of morphogenesis in the vegetative shoot. Development 120, 405–413 (1994).

14. M. Kitagawa, D. Jackson, Plasmodesmata-Mediated Cell-to-Cell Communication in the Shoot Apical Meristem: How Stem Cells Talk. Plants 6, 12 (2017).

15. J. W. Wang, H. J. Squire, N. S. Goh, M. N. Ni, E. Lien, C. Wong, E. González-Grandío, M. P. Landry, Delivered complementation in planta (DCIP) enables measurement of peptide-mediated protein delivery efficiency in plants. Communications Biology 6, 840 (2023).

16. D. Kamiyama, S. Sekine, B. Barsi-Rhyne, J. Hu, B. Chen, L. A. Gilbert, H. Ishikawa, M. D. Leonetti, W. F. Marshall, J. S. Weissman, B. Huang, Versatile protein tagging in cells with split fluorescent protein. Nat Commun 7, 11046 (2016).

17. The UniProt Consortium, A. Bateman, M.-J. Martin, S. Orchard, M. Magrane, S. Ahmad, E. Alpi, E. H. Bowler-Barnett, R. Britto, H. Bye-A-Jee, A. Cukura, P. Denny, T. Dogan, T. Ebenezer, J. Fan, P. Garmiri, L. J. Da Costa Gonzales, E. Hatton-Ellis, A. Hussein, A. Ignatchenko, G. Insana, R. Ishtiaq, V. Joshi, D. Jyothi, S. Kandasaamy, A. Lock, A. Luciani, M. Lugaric, J. Luo, Y. Lussi, A. MacDougall, F. Madeira, M. Mahmoudy, A. Mishra, K. Moulang, A. Nightingale, S. Pundir, G. Qi, S. Raj, P. Raposo, D. L. Rice, R. Saidi, R. Santos, E. Speretta, J. Stephenson, P. Totoo, E. Turner, N. Tyagi, P. Vasudev, K. Warner, X. Watkins, R. Zaru, H. Zellner, A. J. Bridge, L. Aimo, G. Argoud-Puy, A. H. Auchincloss, K. B. Axelsen, P. Bansal, D. Baratin, T. M. Batista Neto, M.-C. Blatter, J. T. Bolleman, E. Boutet, L. Breuza, B. C. Gil, C. Casals-Casas, K. C. Echioukh, E. Coudert, B. Cuche, E. De Castro, A. Estreicher, M. L. Famiglietti, M. Feuermann, E. Gasteiger, P. Gaudet, S. Gehant, V. Gerritsen, A. Gos, N. Gruaz, C. Hulo, N. Hyka-Nouspikel, F. Jungo, A. Kerhornou, P. Le Mercier, D. Lieberherr, P. Masson, A. Morgat, V. Muthukrishnan, S. Paesano, I. Pedruzzi, S. Pilbout, L. Pourcel, S. Poux, M. Pozzato, M. Pruess, N. Redaschi, C. Rivoire, C. J. A. Sigrist, K. Sonesson, S. Sundaram, C. H. Wu, C. N. Arighi, L. Arminski, C. Chen, Y. Chen, H. Huang, K. Laiho, P. McGarvey, D. A. Natale, K. Ross, C. R. Vinayaka, Q. Wang, Y. Wang, J. Zhang, UniProt: the Universal Protein Knowledgebase in 2023. Nucleic Acids Research 51, D523–D531 (2023).

18. K. Mukherjee, L. Brocchieri, T. R. Burglin, A Comprehensive Classification and Evolutionary Analysis of Plant Homeobox Genes. Molecular Biology and Evolution 26, 2775–2794 (2009).

19. M. Zhou, C. Coruh, G. Xu, L. M. Martins, C. Bourbousse, A. Lambolez, J. A. Law, The CLASSY family controls tissue-specific DNA methylation patterns in Arabidopsis. Nat Commun 13, 244 (2022).

20. J.-Y. Kim, Y. Rim, J. Wang, D. Jackson, A novel cell-to-cell trafficking assay indicates that the KNOX homeodomain is necessary and sufficient for intercellular protein and mRNA trafficking. Genes Dev. 19, 788–793 (2005).

21. G. Drin, M. Mazel, P. Clair, D. Mathieu, M. Kaczorek, J. Temsamani, Physico-chemical requirements for cellular uptake of pAntp peptide: Role of lipid-binding affinity. European Journal of Biochemistry 268, 1304–1314 (2001).

22. W. B. Kauffman, T. Fuselier, J. He, W. C. Wimley, Mechanism Matters: A Taxonomy of Cell Penetrating Peptides. Trends in Biochemical Sciences 40, 749–764 (2015).

23. V. Bandmann, J. D. Müller, T. Köhler, U. Homann, Uptake of fluorescent nano beads into BY2-cells involves clathrin-dependent and clathrin-independent endocytosis. FEBS Letters 586, 3626–3632 (2012).

24. K. Aregawi, J. Shen, G. Pierroz, M. K. Sharma, J. Dahlberg, J. Owiti, P. G. Lemaux, Morphogene-assisted transformation of Sorghum bicolor allows more efficient genome editing. Plant Biotechnology Journal 20, 748–760 (2022).

25. P. Agrawal, S. Bhalla, S. S. Usmani, S. Singh, K. Chaudhary, G. P. S. Raghava, A. Gautam, CPPsite 2.0: a repository of experimentally validated cell-penetrating peptides. Nucleic Acids Res 44, D1098–D1103 (2016).

26. K. Numata, Y. Horii, K. Oikawa, Y. Miyagi, T. Demura, M. Ohtani, Library screening of cell-penetrating peptide for BY-2 cells, leaves of Arabidopsis, tobacco, tomato, poplar, and rice callus. Sci Rep 8, 10966 (2018).

27. H. J. Squire, S. Tomatz, J. W.-T. Wang, E. González-Grandío, M. P. Landry, Best Practices and Pitfalls in Developing Nanomaterial Delivery Tools for Plants. ACS Nano 19, 7–12 (2025).

28. S. Chatterjee, E. Kon, P. Sharma, D. Peer, Endosomal escape: A bottleneck for LNP-mediated therapeutics. Proc. Natl. Acad. Sci. U.S.A. 121, e2307800120 (2024).

29. A. Joliot, A. Maizel, D. Rosenberg, A. Trembleau, S. Dupas, M. Volovitch, A. Prochiantz, Identification of a signal sequence necessary for the unconventional secretion of Engrailed homeoprotein. Current Biology 8, 856–863 (1998).

30. E. Dupont, A. Prochiantz, A. Joliot, Identification of a Signal Peptide for Unconventional Secretion. Journal of Biological Chemistry 282, 8994–9000 (2007).

31. H. Chen, D. Jackson, J.-Y. Kim, Identification of evolutionarily conserved amino acid residues in homeodomain of KNOX proteins for intercellular trafficking. Plant Signaling & Behavior 9, e28355 (2014).

32. G. Daum, A. Medzihradszky, T. Suzaki, J. U. Lohmann, A mechanistic framework for noncell autonomous stem cell induction in Arabidopsis. Proc. Natl. Acad. Sci. U.S.A. 111, 14619–14624 (2014).

33. W. J. Lucas, S. Bouché-Pillon, D. P. Jackson, L. Nguyen, L. Baker, B. Ding, S. Hake, Selective Trafficking of KNOTTED1 Homeodomain Protein and Its mRNA Through Plasmodesmata. Science 270, 1980–1983 (1995).

34. Y. Rim, J.-H. Jung, H. Chu, W. K. Cho, S.-W. Kim, J. C. Hong, D. Jackson, R. Datla, J.-Y. Kim, A non-cell-autonomous mechanism for the control of plant architecture and epidermal differentiation involves intercellular trafficking of BREVIPEDICELLUS protein. Functional Plant Biol. 36, 280 (2009).

35. R. K. Yadav, M. Perales, J. Gruel, T. Girke, H. Jönsson, G. V. Reddy, WUSCHEL protein movement mediates stem cell homeostasis in the Arabidopsis shoot apex. Genes Dev. 25, 2025–2030 (2011).

36. J.-Y. Lee, J. Colinas, J. Y. Wang, D. Mace, U. Ohler, P. N. Benfey, Transcriptional and posttranscriptional regulation of transcription factor expression in Arabidopsis roots. Proc. Natl. Acad. Sci. U.S.A. 103, 6055–6060 (2006).

37. R. Shimizu, J. Ji, E. Kelsey, K. Ohtsu, P. S. Schnable, M. J. Scanlon, Tissue Specificity and Evolution of Meristematic WOX3 Function. Plant Physiology 149, 841–850 (2009).

