## Supplementary Materials for "The third alpha helix of plant homeoproteins are generally cell-penetrating to plant cells"

All peptides were chemically synthesized by Lifetein LLC at ≥95% purity with C-terminal amination. Fusion peptides contained a GSGS linker in between GFP11 (N-terminal) and the 3^rd^ alpha helix of interest.

**Methods**

Homeoprotein library construction and 3^rd^ alpha helix identification

Plant homeoproteins (including algae) were mined from the UniProt KB database using the search term: ("homeobox" OR "homeodomain") AND ((taxonomy_id:33090) OR (taxonomy_id:3027) OR (taxonomy_id:38254) OR (taxonomy_id:2763)). Given the database is updated every 8 weeks, throughout the duration of this work, we periodically re-consulted the database with the final search taking place on March 25^th^ 2025. The search on this date yielded 242,886 protein entries, the raw output from this search can be found in data S1. We assumed many of the proteins returned from our search were not actually homeoproteins. We applied the following criteria to filter the list further, first, given the homeodomain is ~60 amino acids in length, any protein entry with less than 60 total amino acids were excluded. Next, if a protein entry did not contain a UniProt annotation for a DNA binding domain or general domain titled “Homeobox” the entry was removed. After filtering, we were left with 48,864 plant homeoprotein entries all of which can be found in data S2. We note 44,681 of these entries contained unique protein sequences.

The UniProt “Homeobox” annotation under the DNA binding domain category and the “Homeobox” annotation under the domain category was used to extract the specific amino acids representing the homeodomain of each protein entry. Of extracted homeodomains, 90% of sequences were of typical with lengths of 55 to 65 amino acids. A small subset was divergent with amino acid lengths longer or shorter lengths than typical homeodomains. Previous work has identified homeodomains with insertions thus for the sake of sequence diversity, we did not exclude these unusual homeodomains. Extracted homedomains for all plant homeoproteins can be found in data S2. Extracted homeodomains were filtered to remove duplicate sequences leaving 20,208 unique homeodomains and then aligned with Clustal Omega. The 3^rd^ alpha helix was identified in the alignment by the highly conserved sequence “WFQN” near the C-terminus of the homeodomain and through reference to the structure of *Arabidopsis thaliana* WUSCHEL as well as the predicted structures of other entries. The aligned homeodomains can be found in data S2. Homeodomains which did not align around this motif were excluded and marked as a sequence with “Poor Alignment”. While the exact length of the 3^rd^ alpha helix domain varies, we choose 17 amino acids centered around the highly conserved tryptophan as representative of the 3^rd^ alpha helix in line with previous work with Penetratin (16 amino acids in length). This resulted in identification of 3,994 unique 3^rd^ alpha helix peptides. The 3^rd^ alpha helixes of all plant homeoproteins can be found in data S2.

Plant growth conditions

*Nicotiana benthamiana* plants were grown in a chamber at 24°C at a light flux of 100 μmol m^-2^ s^-1^ under 16 hours of light and 8 hours of dark. Seeds were sown in square 2.5 inch pots in Lambert LM-HP soil, covered in clear plastic wrap to maintain humidity, and germinated for a week after which individual seedlings were transferred to 2.5 inch pots. Plants were watered weekly supplemented with 20-20-20 fertilizer at a concentration of 75 ppm nitrogen and calcium nitrate at a concentration of 90 ppm nitrogen. *Nicotiana benthamiana* were grown to 4-weeks prior to experimentation.

Transgenic *Arabidopsis thaliana* (Col-0) seedlings expressing the DCIP reporter system were grown in a chamber at 24°C at a light flux of 100 μmol m^-2^ s^-1^ under 16 hours of light and 8 hours of dark. Seeds were sterilized by a 30-second incubation in 70% ethanol followed by a 15-minute incubation in 50-50 water-bleach with <0.5% Tween-20 and rinsed 5x times with sterile water prior to sowing. Seeds were sown in on tissue culture plates in 0.5 mL of sterile half-concentration Murashige Skoog medium supplemented with 1% sucrose (g/mL) at a density of 3-4 seeds per well. Seeds were stratified in the dark at 4°C for 3 days prior to transferring culture plates to growth chambers. Seedlings were grown for 10 days prior to experimentation.

DCIP experiments

DCIP experiments were conducted in five-week old *Nicotiana benthamiana* plants through transient agro-transfection. *Agrobacterium tumefaciens* strain GV3101 bearing the DCIP reporter was grown in LB media supplemented with 50 μg/mL rifampicin, 25 μg/mL gentamycin, and 50 μg/mL kanamycin at 30°C overnight. The overnight cultures were pelleted by centrifugation at 3200 x g for 10 minutes, re-suspended in infiltration buffer (10 mM MES (pH 5.7) and 10 mM MgCl_2_ supplemented with 200 μM acetosyringone), and incubated at room temperature with shaking for four hours. After incubation, the third or fourth mature leaf of *Nicotiana benthamiana* plants were infiltrated on the abaxial side with the buffer culture (OD_600_ of 0.5) using blunt syringes. Three days after infiltration, leaves were infiltrated with peptide dissolved in sterile water using blunt syringes. Infiltration spots were punched with an 8 mm punch and plated on half-concentration Murashige and Skoog medium plates with agar (pH 5.7). Five hours after infiltration with peptide, leaf punches were imaged on a Zeiss LSM880 laser scanning confocal microscope. Images were collected using a Fluar 5x/0.25 M27 air objective with the aperture set to 1 Airy-units on the mCherry channel. Images of sfGFP, mCherry, and chloroplast autofluorescence were collected by excitation with a 488, 561, and 635 nm laser respectively and collection of emission bands at 493-550, 578-645, and 652-728 nm respectively. Between 4 to 12 z-plane images were collected depending on the relative flatness of the field of view.

DCIP experiments were conducted in 10-day old transgenic *Arabidopsis thaliana (Col-0)* seedlings expressing the DCIP reporter system. Tissue culture wells containing seedlings were flooded with 0.5 mL of peptide at the appropriate concentration. Five hours after flooding, seedlings were extracted from media and leaves excised for imaging with a Zeiss LSM880 laser scanning confocal microscope. Images were collected using a Plan-Apochromat 20x/0.8 M27 air objective with the aperture set to 0.75 Airy-units on the mCherry channel. Images of sfGFP, mCherry, and chloroplast autofluorescence were collected by excitation with a 488, 561, and 635 nm laser respectively and collection of emission bands at 493-550, 578-645, and 652-728 nm respectively. Between 4 to 12 z-plane images were collected depending on the relative flatness of the field of view.

Image analysis for DCIP experiments was conducted with Cell Profiler 3.0. For a given z-plane, nuclei and chloroplasts were first segmented with Otsu’s method using the mCherry and chloroplast autofluorescence channels. The segmented chloroplast signal was subtracted from the sfGFP channel and the segmented mCherry nuclei were used as a mask for the sfGFP channel to exclude regions where sfGFP signal did not localize with mCherry nuclei. Maximum intensity projections were constructed from the processed z-stack at a given field of view. Nuclei from maximum intensity projections were filtered for relative circularity to exclude debris. Next, the ratio of the sfGFP intensity to the mCherry intensity (green to red ratio) of each filtered nuclei in a maximum intensity was quantified. To programmatically evaluate if a given nucleus is sfGFP positive, a threshold green to red ratio is set from nuclei of mock treated control leaf discs. For *N. benthamiana* leaves, a nucleus must have a green to red fluorescence ratio greater than 99% of mock nuclei to be considered positive for sfGFP (average green to red ratio of mock nuclei plus 2.57 times the standard deviation of the green to ratio of mock nuclei). For *A. thaliana* seedlings, the average green to red fluorescence ratio of mock treated nuclei was about twice as high that of *N. benthamiana* mock treated nuclei. Thus, to limit false positives, a *A. thaliana* nucleus must have a green to red fluorescence ratio greater than 99.9% of mock treated nuclei (average green to red ratio of mock nuclei plus 3.29 times the standard deviation of the green to ratio of mock nuclei). Individual thresholds were set each for *N. benthamiana* plant to normalize the impact of DCIP reporter expression. In contrast, for *A. thaliana* seedlings an average threshold across all mock treated seedlings was set because individual seedlings cannot be treated with multiple treatments (ie both mock and 3^rd^ alpha helix treated). After identifying the threshold, the number of sfGFP positive nuclei were counted and divided by the total number of nuclei (derived from the mCherry channel) yielding the % GFP positive nuclei which serves as a proxy for delivery efficiency on a cellular basis.

Cre delivery experiments

Cre delivery experiments were conducted in five-week old *Nicotiana benthamiana* plants through transient agro-transfection. *Agrobacterium tumefaciens* strain GV3101 bearing the Cre reporter system shown in Figure 4 A were grown in LB media supplemented with 50 μg/mL rifampicin, 25 μg/mL gentamycin, and 50 μg/mL spectinomycin at 30°C overnight. The overnight cultures were pelleted by centrifugation at 3200 x g for 10 minutes, re-suspended in infiltration buffer (10 mM MES (pH 5.7) and 10 mM MgCl_2_ supplemented with 200 μM acetosyringone), and incubated at room temperature with shaking for four hours. After incubation, the third or fourth mature leaf of *Nicotiana benthamiana* plants were infiltrated on the abaxial side with the buffer culture (OD_600_ of 0.5) using blunt syringes. Two days after infiltration, leaves were infiltrated with 10 µM of either recombinant Cre-ClWOX or recombinant Cre with no 3^rd^ alpha helix tag using blunt syringes. After 24 hours, infiltration spots were punched with an 8 mm punch and and imaged on a Zeiss LSM880 laser scanning confocal microscope. Images were collected using a Fluar 5x/0.25 M27 air objective or Plan-Apochromat 20x/0.8 M27 air objective with the aperture set to 1 Airy-units on the mCherry channel. Images of GFP, mCherry, and chloroplast autofluorescence were collected by excitation with a 488, 561, and 635 nm laser respectively and collection of emission bands at 493-550, 578-645, and 652-728 nm respectively. Between 4 to 12 z-plane images were collected depending on the relative flatness of the field of view.

Cell Profiler 3.0 was used to programmatically quantify delivery events. For a given z-plane, nuclei and chloroplasts were first segmented with Otsu’s method using mCherry and chloroplast autofluorescence channels. The segmented chloroplast signal was subtracted from the GFP channel and the segmented mCherry nuclei were used as a mask for the GFP channel to exclude areas where GFP signal did not localize with mCherry nuclei. Maximum intensity projections were constructed from the processed z-stack at a given field of view. Nuclei from maximum intensity projections were filtered for relative circularity to exclude debris and the ratio of GFP intensity to mCherry intensity (green to red ratio) of each filtered nuclei was quantified. GFP positive nuclei were identified by green to red intensity ratio which was greater than 99% of nuclei from mock treated control leaves.

Plasmids

Plasmids for *Agrobacterium tumefaciens* transfection were built using GoldenBraid 2.0 (GB2.0) modular cloning system. All plasmid cloning was conducted in XL1-blue *E. coli* strains. The previously described DCIP plasmid was assembled into an alpha1R backbone per the standard GB2.0 protocol and can be found on addgene (Plasmid #193860) (*1*). The Cre reporter system shown in Figure 4 A was constructed by first assembling the mCherry transcriptional unit into an alpha1R backbone per the standard GB2.0 protocol with 35s promoter and Nos terminator. The GFP transcriptional unit was assembled into an alpha2R by adding flanking loxP sites to a Nos terminator via PCR 5’ of the reading frame of GFP with 35s promotor and Nos terminator. The two transcriptional units were combined into a single omega1 backbone per the standard GB2.0 protocol.

Recombinant protein expression and purification

Recombinant proteins were expressed in freshly transformed BL21 (DE3) *E. coli.* Single colonies were seeded into 2 mL of LB supplemented with 50 μg/mL kanamycin and shaken overnight at 37°C. Overnight starter cultures were transferred (800 μL) to 50 mL of LB (no antibiotics) and grown at 37°C to an OD_600_ of 0.4-0.6 prior to cooling to 18°C. Cooled cultures were induced with 0.5 mM IPTG and shaken for 16 hours at 18°C. Cultures were pelleted, decanted, and stored at -80°C for future purification.

Recombinant 6x His tagged proteins were purified using nickel ion affinity chromatography. A pellet from a 50 mL culture was suspended in 5 mL of lysis buffer (100 mM NaH2PO4, 10 mM Tris, 300 mM NaCl, 10 mM imidazole pH 8.0 with protease inhibitor cocktail) and probe tip sonicated for five minutes at 50% amplitude with 1 second pulses (on/off). Lysed culture was pelleted at 38k x g for 30 minutes at 4°C and decanted. The soluble fraction was incubated with 1 mL of HisPur Ni NTA slurry at 4°C for 45 minutes with gentle shaking. After incubation, the slurry was washed in batch by centrifuging for 2 minutes at 700 x g and removing the supernatant. The slurry was first washed with 5 mL of lysis buffer followed by 10 mL of lysis buffer at 1 M NaCl. Protein was eluted in batch 1 mL at a time with lysis buffer at 250 mM imidazole for a total of four elutions. Proteins were dialyzed against 33 mM NaCl, 10 mM MgCl_2_, 50 mM Tris-HCl pH 8.0 for 3 hours in 3.5 kDa cut-off membranes.

In-Vitro GFP11 complementation assay

Recombinant sfGFP1-10 (10 mM Tris, 10 mM NaCl pH 7.4) at 10 µM and peptide at 20 µM was combined in equal volumes to a final volume of 10 µL in 96-well PCR plates. Each peptide was run in triplicate. Complementation was tracked overtime using a Biorad CFX96 and reported at 5 hours post-mixing. To normalize across plates, the complementation fluorescence of all peptides was normalized to the complementation fluorescence of GFP11 alone with no linker or 3^rd^ alpha helix tag.

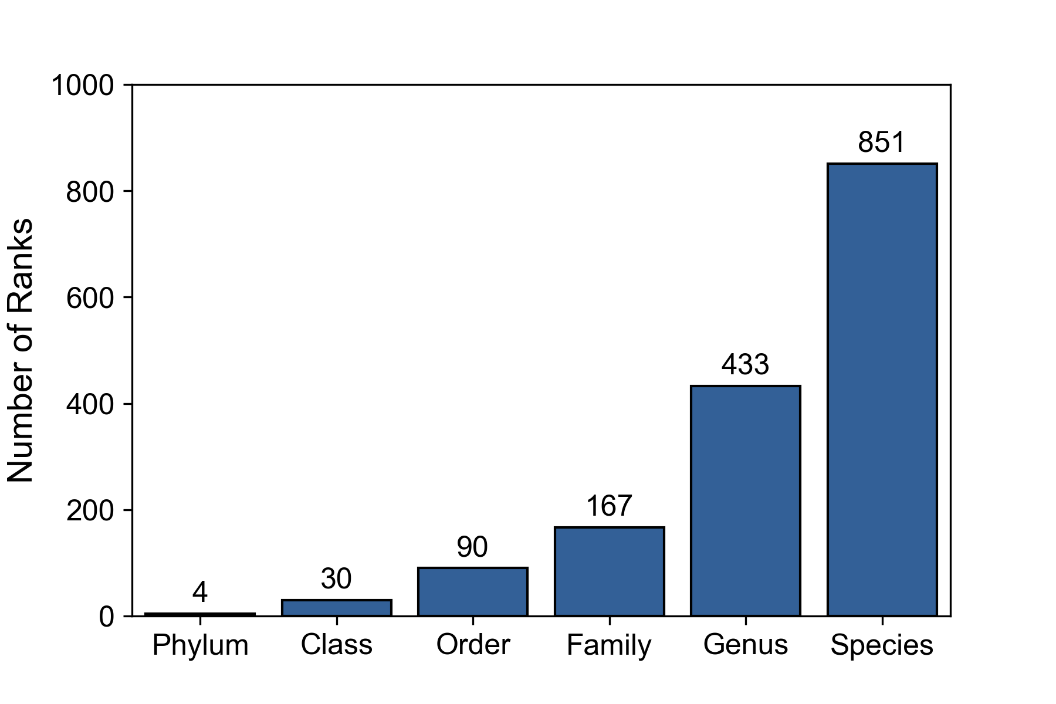

**Fig. S1. Species diversity of identified plant homeoproteins.**

Bar graph of the number of unique ranks found at each taxonomic level for the 40,351 plant homeoproteins identified on UniProt.

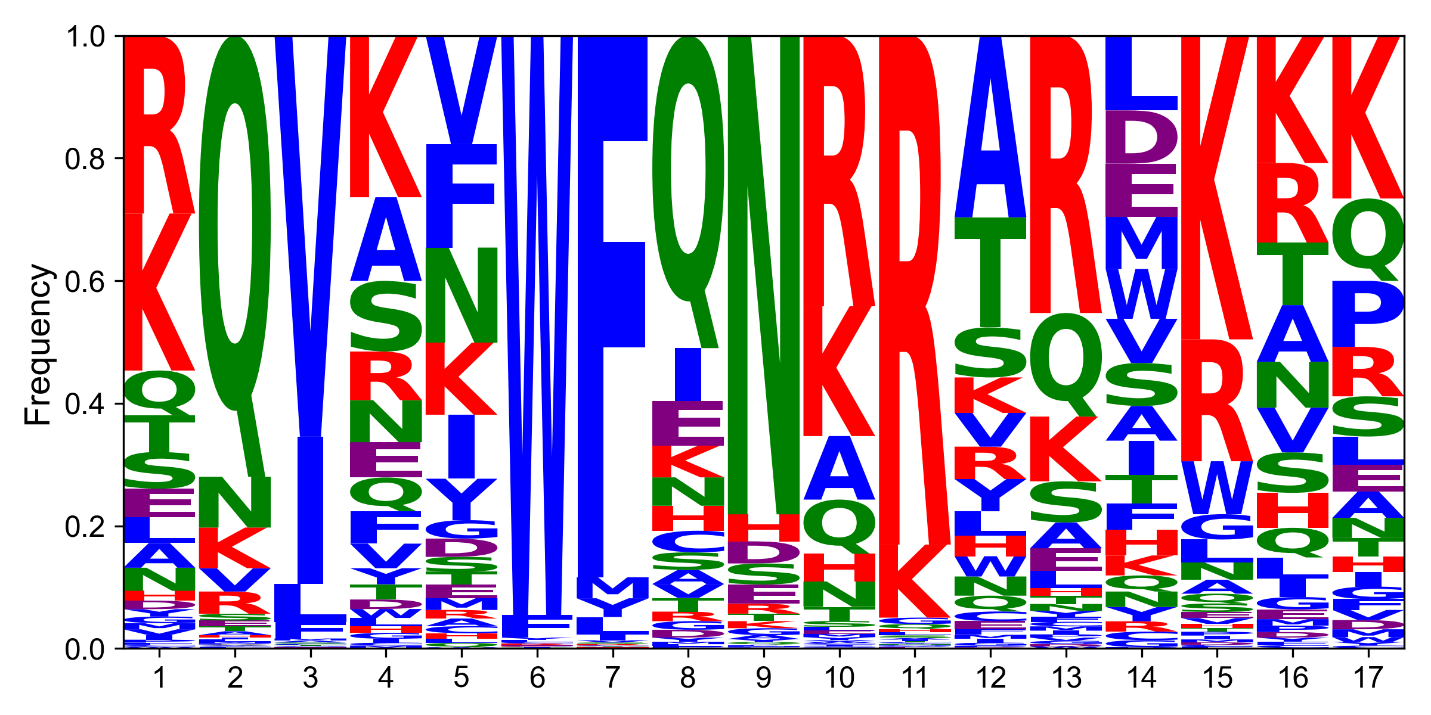

**Fig S2. Plant 3^rd^ alpha helix peptide logo sequence.**

Amino acid frequency logo by position for all 3,994 unique plant 3^rd^ alpha helix peptides cataloged. Residues are colored based off chemical similarity blue = hydrophobic (A, C, I, L, M, F, W, V, P, Y, G), green = polar uncharged (N, Q, S, T), purple = negatively charged (D, E), and red = positively charged (H, K, R).

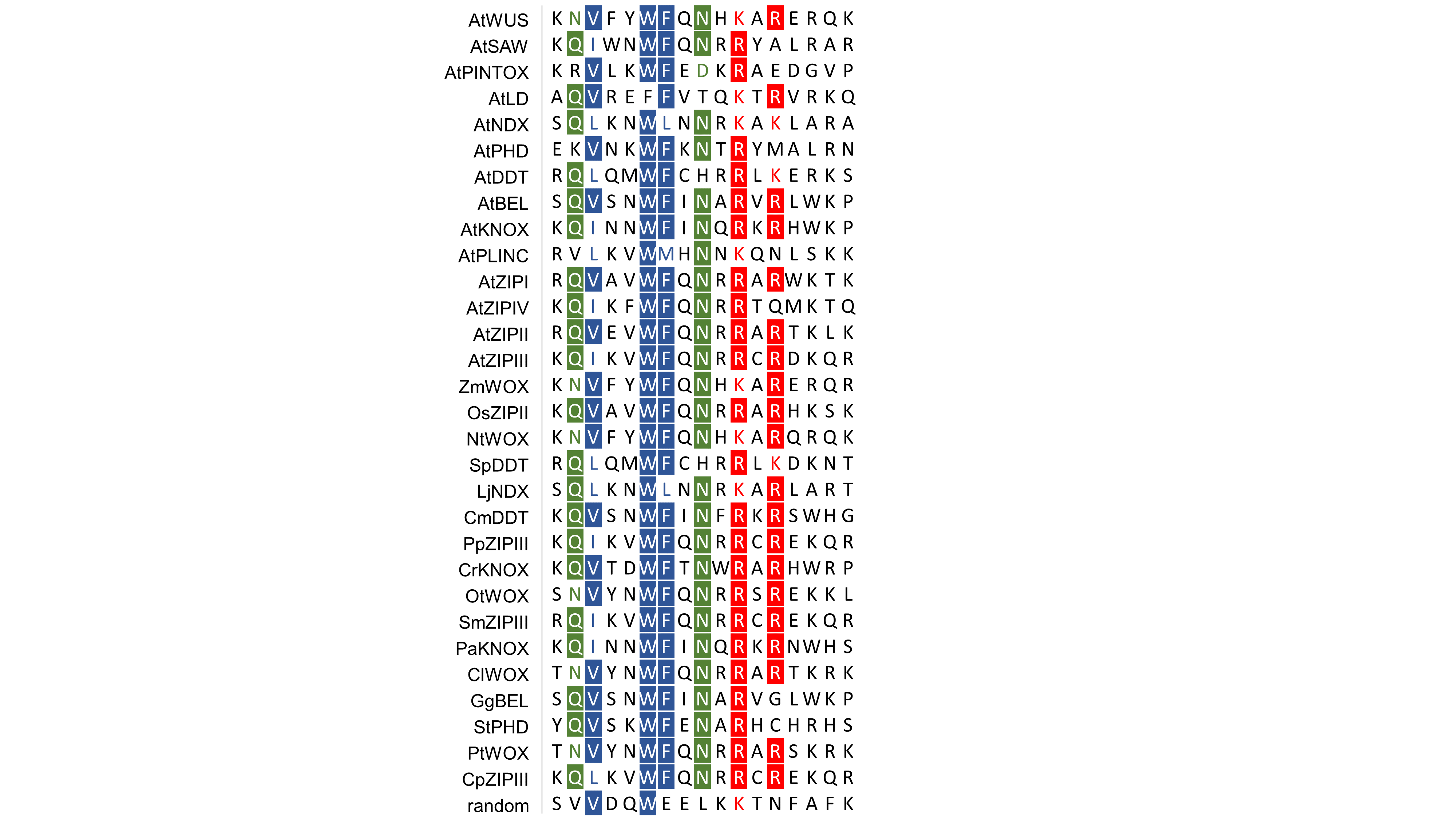

**Fig. S3. Alignment of experimentally tested plant 3^rd^ alpha helix peptides.**

Sequence alignment of the 30 plant 3^rd^ alpha helix peptides selected for experimentation as well as one non-homeoprotein derived sequence (labeled random). Positions with 60% or more conservation of the same residue are colored based off chemistry. At these positions, the most conserved residue is boxed in color with white lettering and resides with chemical similarity are marked with colored lettering. Residues are colored based off chemical similarity blue = hydrophobic (A, C, I, L, M, F, W, V, P, Y, G), green = polar uncharged (N, Q, S, T), purple = negatively charged (D, E), and red = positively charged (H, K, R).

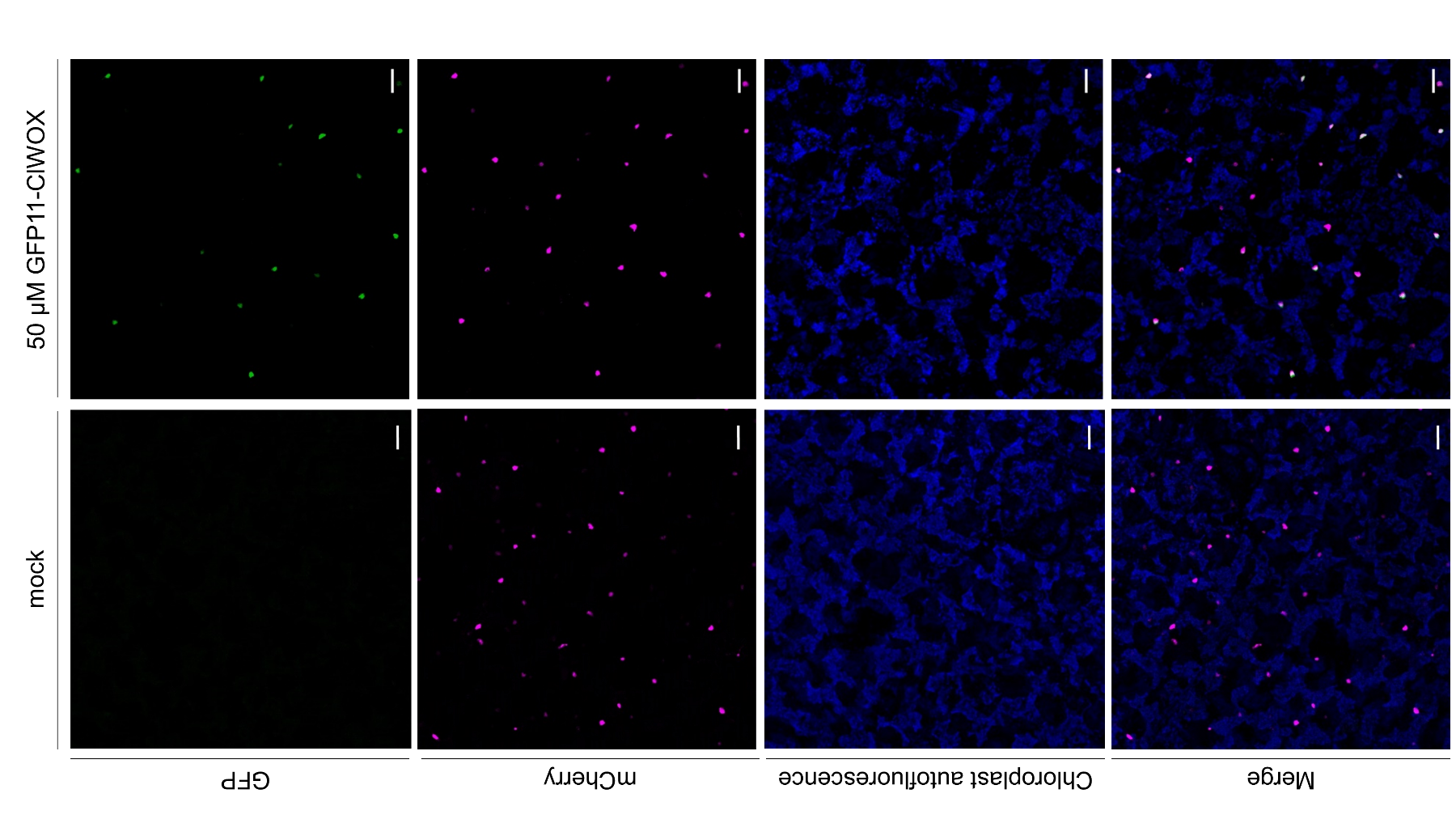

**Fig S4. DCIP confocal slice with chloroplast overlay.**

Representative single slice confocal micrographs of *N. benthamiana* leaves expressing the DCIP reporter, mock treated or treated with 50 µM of ClWOX fused to GFP11. Scale bar 50 µm.

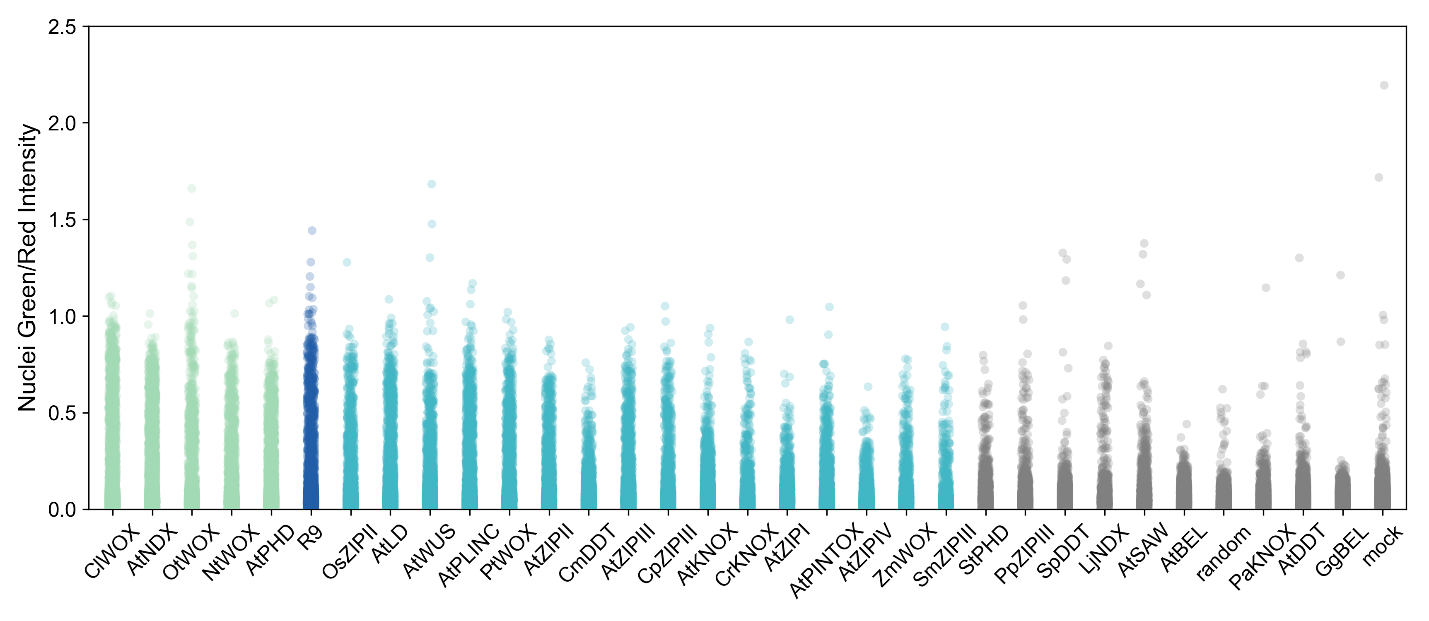

**Fig. S5. Ratio of green to red nuclear intensities in DCIP expressing *N. benthamiana leaves.***

Green to red intensity for every *N. benthamiana* nucleus screened in Fig. 2B for each 3^rd^ alpha helix peptide. Between 1,800 and 4,500 nuclei were screened across 6 biological replicates per 3^rd^ alpha helix peptide. Green to red intensities were used to classify nuclei as GFP positive or GFP negative to determine cellular internalization efficiencies.

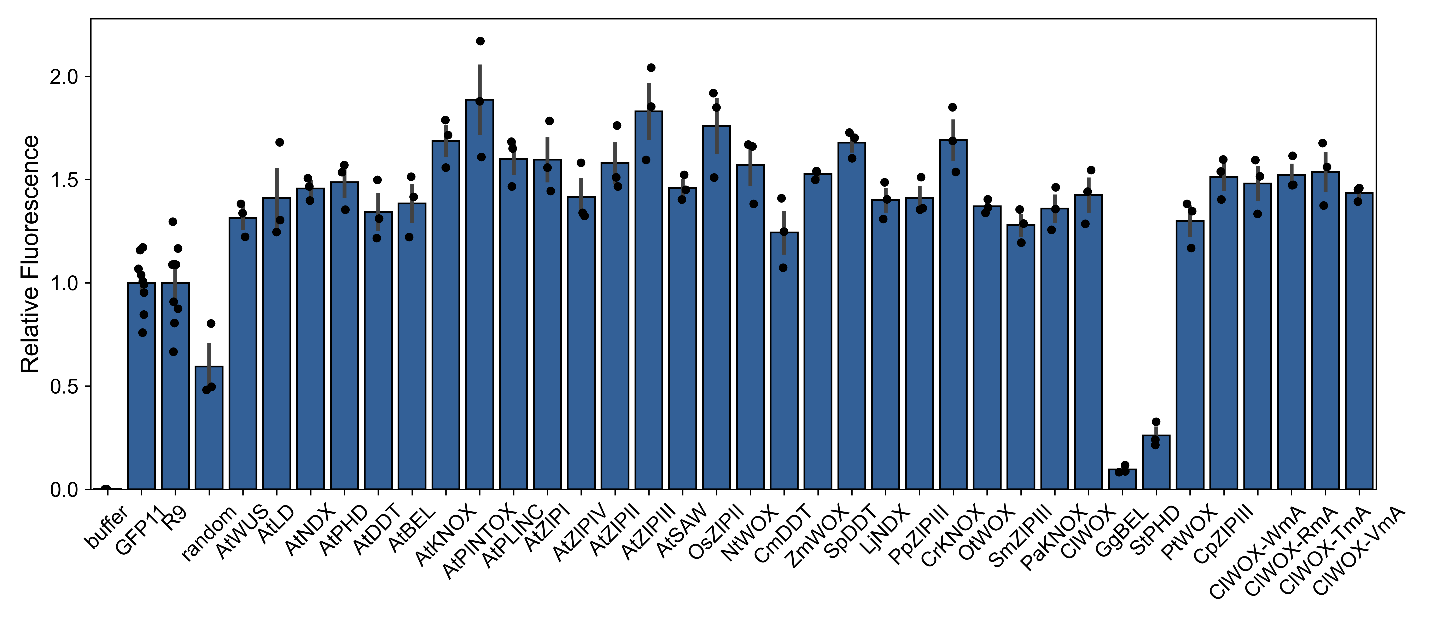

**Fig. S6.** ***In-vitro* fluorescence complementation.**

*In-vitro* complementation assay for different fusion peptides containing GFP11. Fluorescence from complementation is measured 5 hours after combining recombinant GFP1-10 and GFP11 fusion peptide.

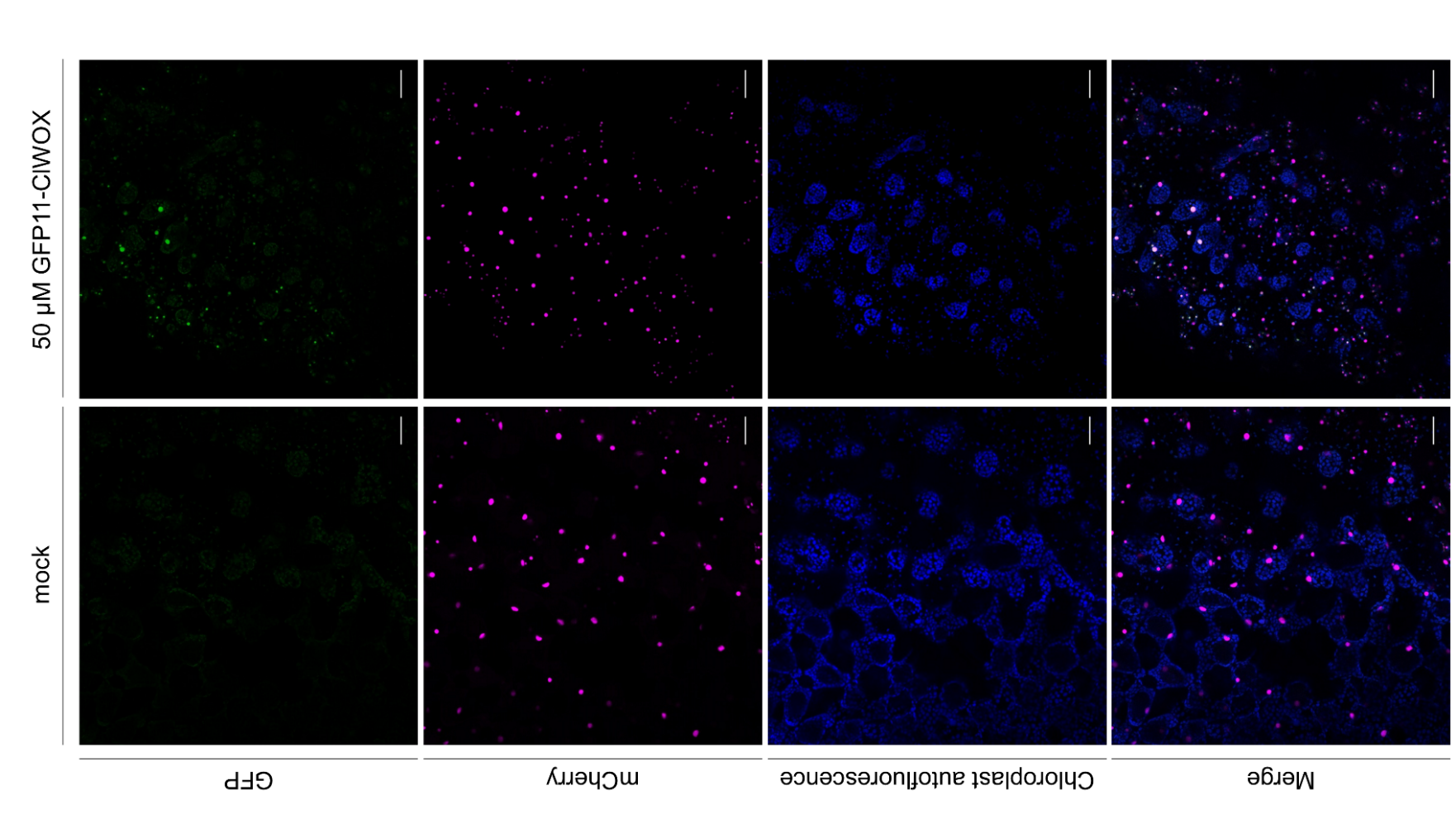

**Fig. S7. Maximum intensity projections of DCIP *A. thaliana* with chloroplast overlay.**

Representative maximum intensity projections of confocal micrographs of the leaves of *A. thaliana* seedlings expressing DCIP mock treated or treated with 50 µM ClWOX fused to GFP11. Scale bars represent 50 µm.

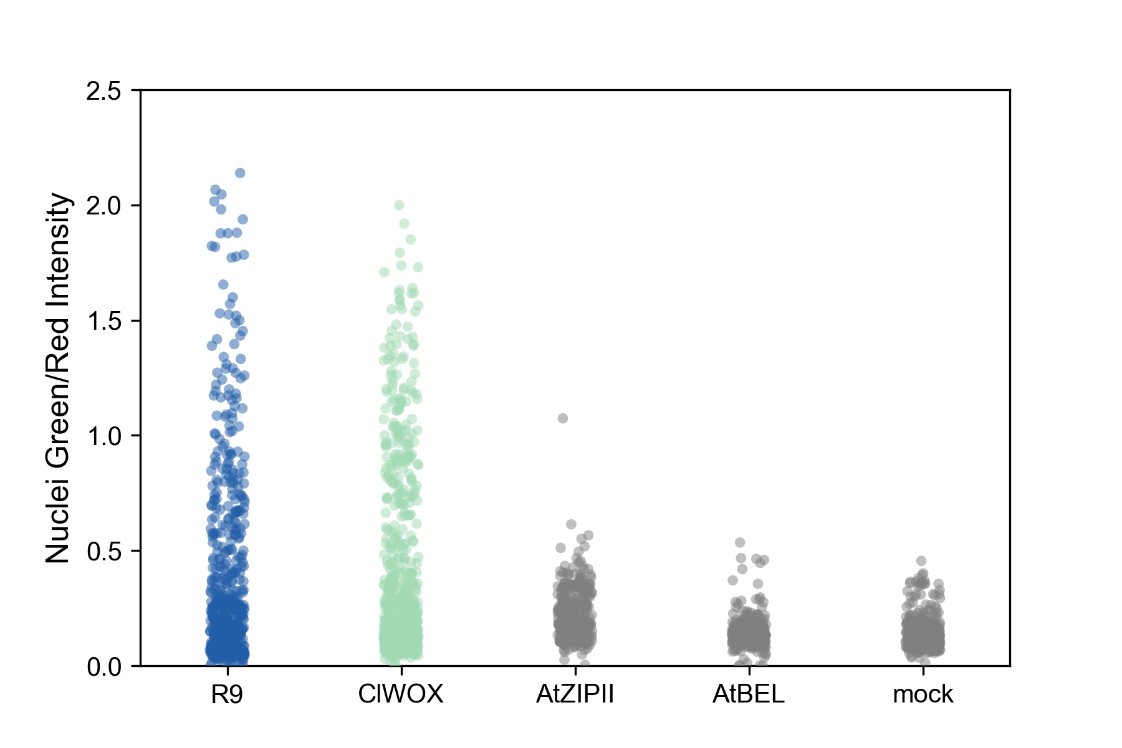

**Fig. S8. Ratio of green to red nuclear intensities in DCIP expressing *A. thaliana.***

Green to red intensity for every *A. thaliana* nucleus screened in Figure 2 D for each 3^rd^ alpha helix peptide. Between 302 and 786 nuclei were screened across 6 biological replicates per 3^rd^ alpha helix peptide. Green to red intensities were used to classify nuclei as GFP positive or GFP negative to determine cellular internalization efficiencies.

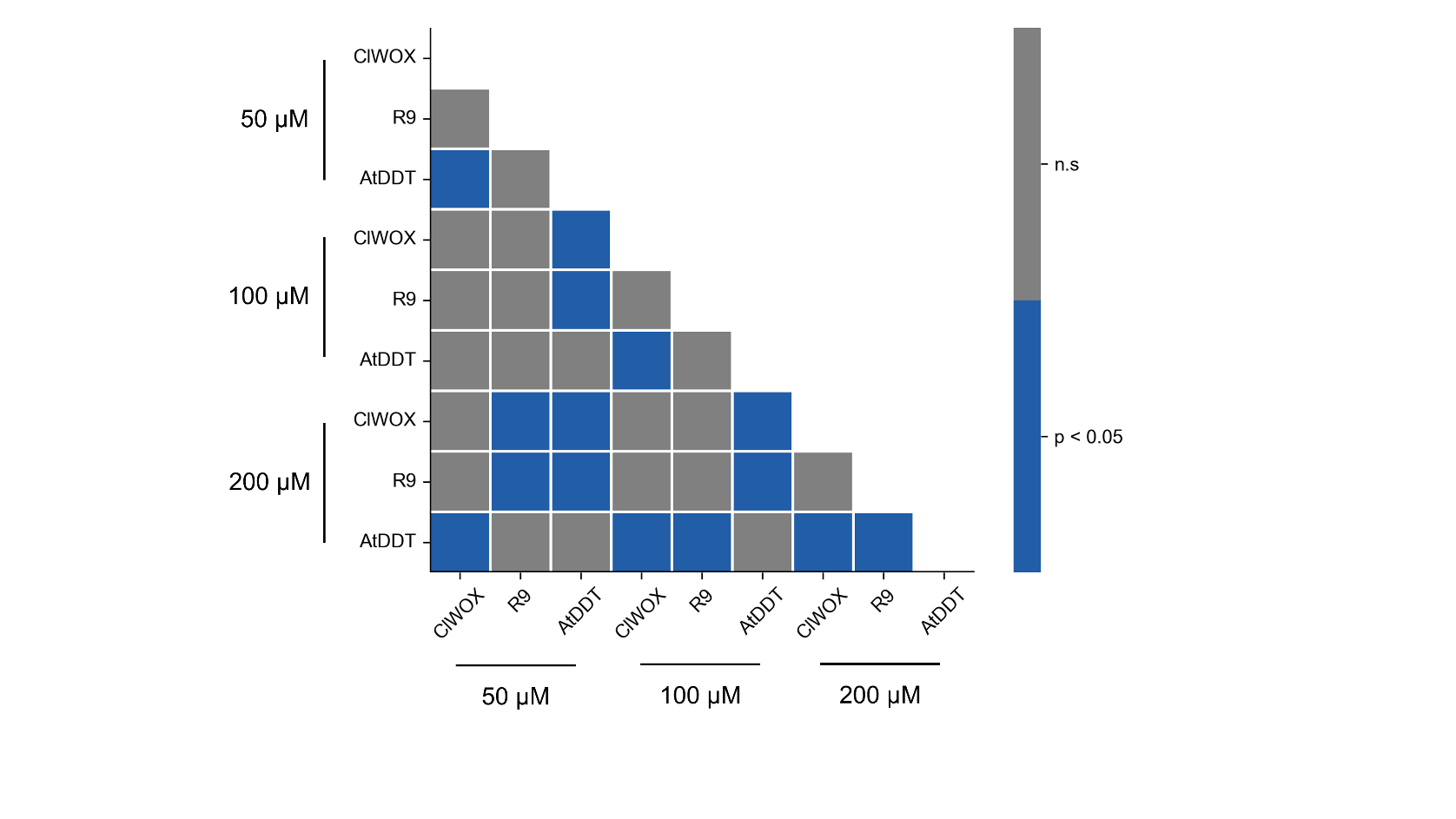

**Fig. S9. Pair-wise comparison of cellular internalization efficiency at different peptide concentrations.**

Results of a Kruskal-Wallis test followed by Dunn’s multiple comparison post-hoc on data in Fig. 3C (*N. benthamiana* leaves expressing DCIP treated with various peptides at various concentrations).

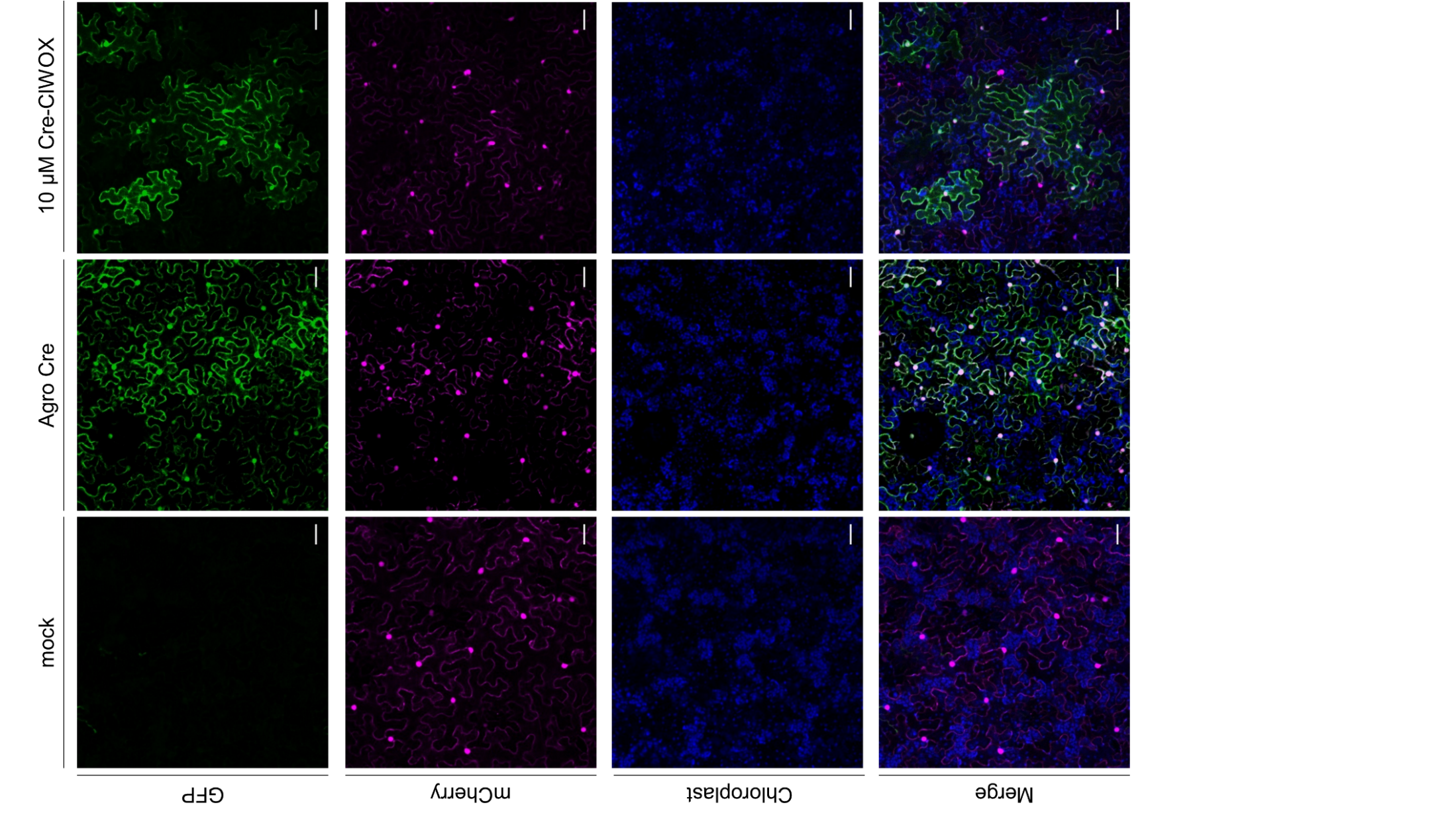

**Fig. S10. Maximum intensity projection of *N. benthamiana* leaves expressing Cre reporter.**

Representative maximum intensity projections of confocal micrographs of *N. benthamiana* leaves expressing a Cre delivery reporter system under, mock treatment, *A. tumefaciens* (bearing Cre plasmid as a positive control) infiltration, and 10 µM Cre-ClWOX infiltration. Scale bars represent 50 µm.

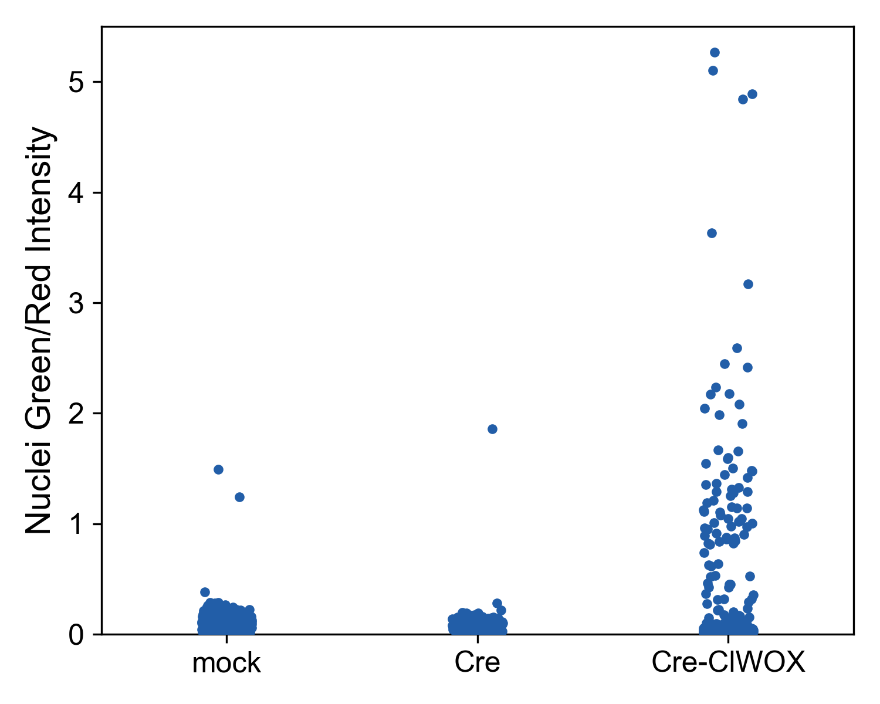

**Fig. S11.** **Ratio of green to red nuclear intensities in Cre reporter expressing *N. benthamiana*.**

Green to red intensity for every *N. benthamiana* nucleus screened in Figure 4B for delivery of Cre recombinase. Between 280 and 582 nuclei were screened across 6 biological replicates. Green to red intensities were used to classify nuclei as GFP positive or GFP negative to determine cellular internalization efficiencies.

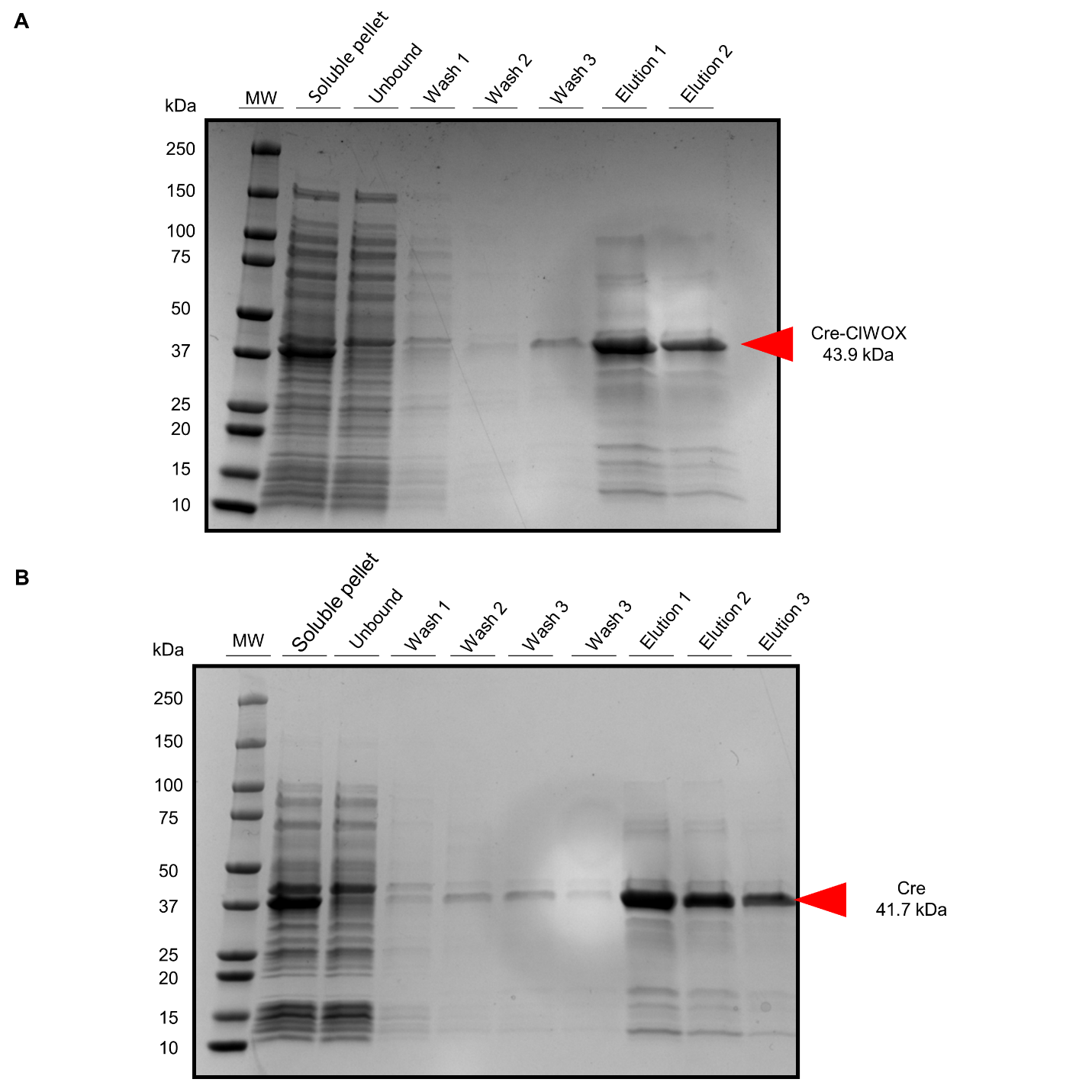

**Figure S12. SDS-PAGE gels for recombinant proteins.**

SDS-PAGE gel for purifications of recombinant Cre-ClWOX (43.9 kDa) and Cre (41.7 kDa).

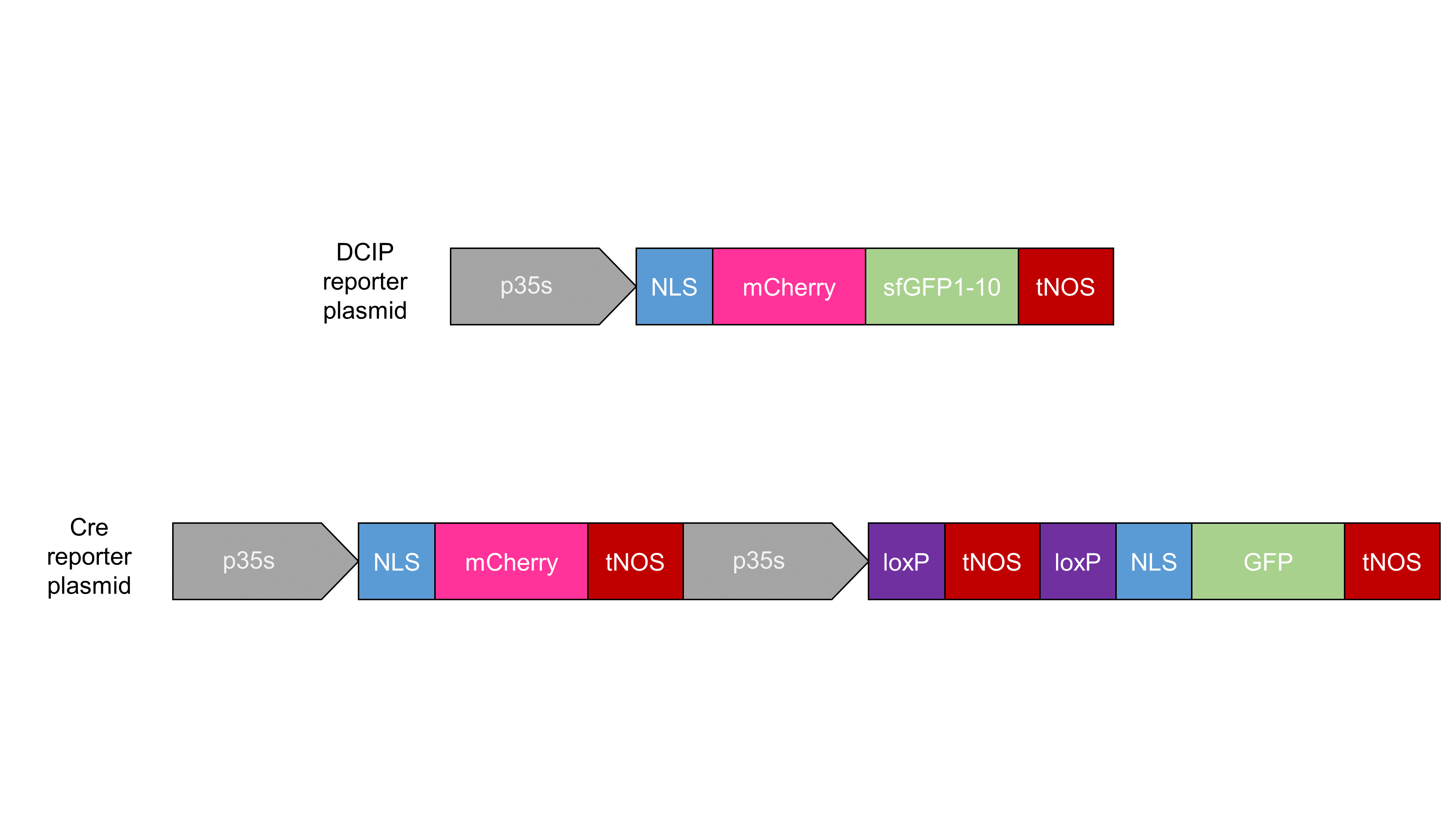

**Fig. S13. Reporter plasmids**.

Schematic of the DCIP and Cre reporter plasmids.

**Table S1. Plant homeodomain trafficking data.**

Literature data on the ability of different of plant homeoproteins to traffic and data on the ability of the corresponding 3^rd^ alpha helix to internalize to plant cells. N/A indicates the specific 3^rd^ alpha helix of the corresponding homeoprotein was not screened for internalization capability.

| Homeodomain Protein | UniProt | Protein Traffics? | 3^rd^ Alpha Helix Internalizes? | Reference |
| --- | --- | --- | --- | --- |
| KNOTTED1 | P24345 | Y | Y | Lucas et al. (*2*) |
| STM | Q38874 | Y | Y | Kim et al. (*3*) |
| KNAT1 | P46639 | Y | Y | Rim et al. (*4*) |
| WUS | Q9SB92 | Y | Y | Yadav et al. (*5*) |
| WOX5 | Q8H1D2 | Y | Y | Daum et al. (*6*) |
| REV | Q9SE43 | Y | Y | Lee et al. (*7*) |
| LeT6 | O22299 | Y | Y | Kim et al. (*3*) |
| KNAT3 | P48000 | N | N | Kim et al. (*3*) |
| BLR | Q9LZM8 | N | N | Kim et al. (*3*) |
| KNAT2 | P46640 | N | Y | Kim et al. (*3*) |
| KNAT6 | Q84JS6 | N | Y | Kim et al. (*3*) |
| NARROWSHEATH1 | Q70UV1 | N | Y | Shimizu et al. (*8*) |
| WOX3 | Q9SIB4 | N | N/A | Shimizu et al. (*8*) |
| WOX13 | O81788 | N | N/A | Yadav et al. (*5*) |

**References**

1.	J. W. Wang, H. J. Squire, N. S. Goh, M. N. Ni, E. Lien, C. Wong, E. González-Grandío, M. P. Landry, Delivered complementation in planta (DCIP) enables measurement of peptide-mediated protein delivery efficiency in plants. *Communications Biology* **6**, 840 (2023).

2.	W. J. Lucas, S. Bouché-Pillon, D. P. Jackson, L. Nguyen, L. Baker, B. Ding, S. Hake, Selective Trafficking of KNOTTED1 Homeodomain Protein and Its mRNA Through Plasmodesmata. *Science* **270**, 1980–1983 (1995).

3.	J.-Y. Kim, Y. Rim, J. Wang, D. Jackson, A novel cell-to-cell trafficking assay indicates that the KNOX homeodomain is necessary and sufficient for intercellular protein and mRNA trafficking. *Genes Dev.* **19**, 788–793 (2005).

4.	Y. Rim, J.-H. Jung, H. Chu, W. K. Cho, S.-W. Kim, J. C. Hong, D. Jackson, R. Datla, J.-Y. Kim, A non-cell-autonomous mechanism for the control of plant architecture and epidermal differentiation involves intercellular trafficking of BREVIPEDICELLUS protein. *Functional Plant Biol.* **36**, 280 (2009).

5.	R. K. Yadav, M. Perales, J. Gruel, T. Girke, H. Jönsson, G. V. Reddy, WUSCHEL protein movement mediates stem cell homeostasis in the *Arabidopsis* shoot apex. *Genes Dev.* **25**, 2025–2030 (2011).

6.	G. Daum, A. Medzihradszky, T. Suzaki, J. U. Lohmann, A mechanistic framework for noncell autonomous stem cell induction in *Arabidopsis*. *Proc. Natl. Acad. Sci. U.S.A.* **111**, 14619–14624 (2014).

7.	J.-Y. Lee, J. Colinas, J. Y. Wang, D. Mace, U. Ohler, P. N. Benfey, Transcriptional and posttranscriptional regulation of transcription factor expression in *Arabidopsis* roots. *Proc. Natl. Acad. Sci. U.S.A.* **103**, 6055–6060 (2006).

8.	R. Shimizu, J. Ji, E. Kelsey, K. Ohtsu, P. S. Schnable, M. J. Scanlon, Tissue Specificity and Evolution of Meristematic WOX3 Function. *Plant Physiology* **149**, 841–850 (2009).
